# The role of ERFVIIs as oxygen-sensing transducers in the evolution of land plant response to hypoxia

**DOI:** 10.1101/2024.09.27.615240

**Authors:** Laura Dalle Carbonare, Hans van Veen, Vinay Shukla, Monica Perri, Liem Bui, Michael J. Holdsworth, Francesco Licausi

## Abstract

The transcriptional response to low oxygen (hypoxia) in the flowering plant *Arabidopsis thaliana* is transduced through group VII Ethylene Response Factor (ERFVII) transcription factors, whose proteolysis is oxygen-dependent via the PLANT CYSTEINE OXIDASE (PCO) N-degron pathway. When and how this response to hypoxia evolved in land plants remains unknown. Here we investigated the conservation and divergence of transcriptional responses to hypoxia in major land plant clades. We identified induction of gene functions associated with glycolysis and fermentation as part of a conserved response across all land plant divisions. Our results indicate that ERFVIIs appeared in the last common ancestor of vascular plants with true roots, concurrently with oxygen-dependent destabilisation, to regulate hypoxia-adaptive genes. Proteins from other ERF groups have been recruited multiple times in different clades as substrates of the PCO N-degron pathway. Our results demonstrate that the response of land plants to hypoxia has been refined in derived clades through the evolution of ERFVIIs as transcriptional transducers, that occurred concomitantly with the appearance of vascular systems and roots as foraging systems through hypoxic soil.

## Introduction

Colonisation of land by plants was one of the most important evolutionary events in the history of our planet, since it shaped Earth’s atmosphere and terrestrial ecology (Morris et al. 2018). This event, which occurred around 480 million years ago, contributed to the stabilisation and increase in molecular oxygen (O_2_) concentration in the atmosphere by the end of the Ordovician period, thanks to the photosynthetic activity of land plants (Donoghue et al. 2021). Fossil records indicate that primordial land plants were phenotypically similar to extant bryophytes and thrived in highly humid habitats such as riparian sites and marshes, often subjected to floods events (Bowman et al. 2017). One of the main perturbations that occurs in plants when subjected to flooding is the reduced diffusion of gases, principally O_2_, within tissues (Voesenek and Bailey-Serres 2015). Response to low oxygen availability (hypoxia) has been extensively studied in flowering plants – including the model species *Arabidopsis thaliana* and *Oryza sativa* - and consists of the concerted action of Group VII Ethylene Response Factor (ERFVII) transcription factors and Plant Cysteine Oxidase (PCO) oxygen-sensing enzymes through the PRT6 N-degron pathway (Licausi et al. 2011; Gibbs et al. 2011; Weits et al. 2014; White et al. 2017). Under normoxia, constitutively expressed ERFVIIs are degraded by virtue of a conserved N-terminal Cys-, generated following removal of Met1 by constitutive Methionine Amino Peptidase (MAP) activity. In the presence of O_2_ and nitric oxide (NO), oxidation of the sulphur in the N-terminally exposed Cys to sulfinic acid in the ERFVII degron by PCO, generates a substrate for subsequent arginylation by Arginyl Transferase (ATE) and subsequent polyubiquitylation via the concerted action of E3 ubiquitin-ligases PROTEOLYSIS 6 (PRT6) and BIG (Zhang et al. 2024), leading to degradation through the proteasome (Zubrycka et al. 2023). PCO activity is impaired under oxygen-limiting conditions and, consequently, stabilised ERFVIIs accumulate in the nucleus where they regulate gene expression leading to plant adaptation to low O_2_ conditions. Since O_2_ is the terminal acceptor of the electron flux in oxidative phosphorylation, one of the primary effects of reduced O_2_ availability in green plants is the metabolic switch from mitochondrial respiration to fermentation to sustain ATP synthesis (Voesenek and Bailey-Serres 2015; van Dongen and Licausi 2015; Bui et al. 2019). Despite the conservation of the enzymatic components of the N-degron pathway in eukaryotes (Hammarlund et al. 2020; Holdsworth and Gibbs 2020) and the importance of low O_2_ sensing for flowering plant adaptation to hypoxia (Xu et al. 2006; Abbas et al. 2022; Weits et al. 2019), little is known about low O_2_ sensing, signalling and molecular responses outside angiosperms. This prevents a comparison of the evolution of plant and animal oxygen sensing mechanisms. Indeed, while recent studies placed the origin of metazoan Hypoxia Inducible Factor (HIF) in the early Cryogenian (Song et al. 2022; Mills et al. 2018), a phylogenetic reconstruction of ERFVIIs as O_2_-dependent N-degron substrates is lacking. Here, we present a comprehensive study of the evolution of the land plant regulation of the transcriptional response to hypoxia. We reconstruct the evolution of the ERFVIIs, their regulation through the PRT6 N-degron pathway and their role as O_2_ sensing transducers in land plants.

## Results

### Transcriptional responses to hypoxia in land plants

We compared the transcriptional response to hypoxia in a range of species that recapitulate the major events in the evolution of land plants. We chose *Marchantia polymorpha* (liverwort) and *Physcomitrium patens* (moss) for bryophytes, *Selaginella moellendorffii* for lycophytes, *Equisetum hyemale* and *Pteris vittata* for eusporangiate and leptosporangiate ferns (**Fig. 1A**). Given the differences in life cycle in each division, we selected the developmental phase that includes the main photosynthetically active organs, to enable comparability of tissues with similar metabolic features. Therefore, we treated gametophores of *M. polymorpha* (n) and *P. patens* (n), and mature sporophytes of *S. moellendorffii* (2n), *E. hyemale* (2n) and *P. vittata* (2n) with 1% ambient O_2_ in the dark for 8h. Controls were maintained under aerobic conditions (21% ambient O_2_) in the dark. Differentially expressed genes (DEGs) were identified through mRNA sequencing (**Supplementary Tables 1-5**). We compared the effect of hypoxia on these five species with hypoxia transcriptome datasets available for photosynthetic sporophytes of angiosperms (*A. thaliana* (Licausi et al. 2011) and *Oryza sativa* (Narsai et al. 2009)). We observed different magnitudes of response for these species, where *P. patens* exhibited the least changes, in terms of both number of DEGs and extent of up- and down-regulation (**Fig. 1B**). In contrast, *A. thaliana* and *O. sativa* showed the strongest response to the treatment (**Fig. 1B**).

**Figure 1.**
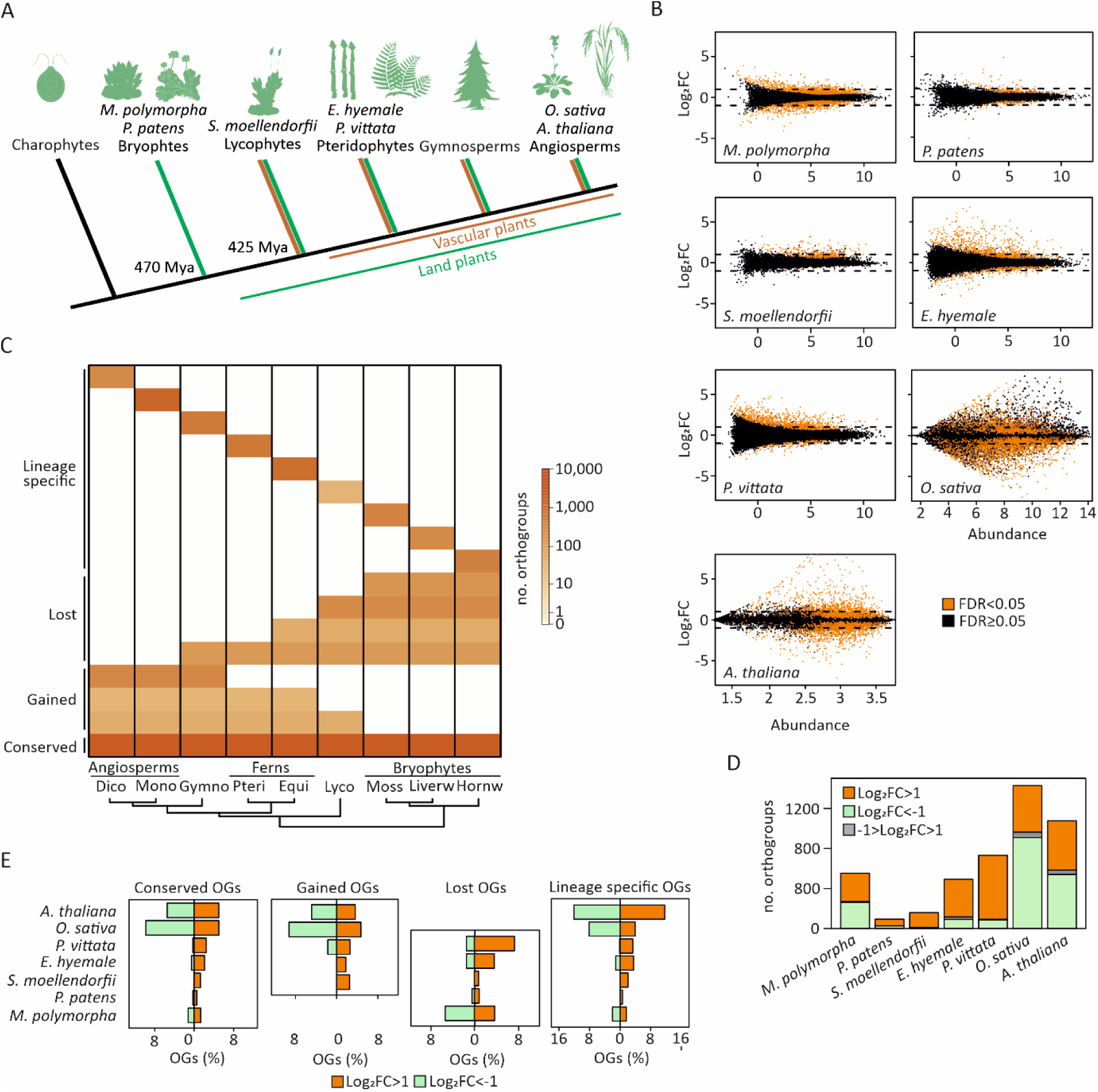
The transcriptional response to hypoxia across land plant divisions. **A**) Current view of the timeline for the divergence of phylogenetic groups. First land plants colonised the land around 470 million years ago (Mya) and possessed features similar to the extant bryophytes, whereas the first vascular plants appeared on land around 425 Mya (Donoghue et al. 2021). **B**) Funnel plots comparing the change genes expression, expressed as log_2_FC, in response to 8 hours treatment with hypoxia (1% v/v O_2_) compared to normoxia of *M. polymorpha*, *P. patens*, *S. moellendorfii*, *E. hyemale*, *P.* vittata, O*. sativa* and *A. thaliana*). **C**) Heatmap representing the number of orthogroups belonging to the ‘conserved’, ‘gained’, ‘lost’, and ‘lineage specific’ categories, for each main group of land plants. **D**). Bar plot showing the number of orthogroups containing upregulated (log_2_FC>1), downregulated (log_2_FC<-1), or simultaneously up- and downregulated (-1> log_2_FC>1) genes for the dataset described in (**B**). **E**) Bar plots showing the percentage (%) of orthogroups up (log_2_FC>1) or down (log_2_FC<-1) regulated by hypoxia (1% v/v O_2_, 8 hours) for the seven species described in (**B**). The orthogroups are classified as ‘conserved’, ‘gained’, ‘lost’ or ‘lineage specific’ (**Supplementary Fig S1-2A**).

The diversification of genes across plant taxa strongly limited an exhaustive analysis of the conservation and divergence of the response to hypoxia between lineages. To aid comparisons, we identified gene sets that originated from a single gene in the last common ancestor of all the species under consideration, termed orthogroups (Wapinski et al. 2007). To define orthogroups, we applied the OrthoFinder method (Emms and Kelly 2015) to the transcriptomes of 33 species corresponding to nine clades: dicot and monocot angiosperms, gymnosperms, leptosporangiate ferns, horsetail ferns, lycophytes, mosses, liverworts and hornworts, including the seven species for which we generated or retrieved hypoxia transcriptomes (**Supplementary Tables 6-12**). We identified 4,969 orthogroups conserved in all clades, and 1,543 and 808 orthogroups that were gained and lost in angiosperms respectively. 10,281 orthogroups were found to be specific to one of the nine clades (**Fig. 1C**, **Supplementary Table 13**, **Supplementary Fig. 1A-B, 2A**). Among conserved orthogroups, our analysis revealed a high number of orthologues identified among more related species (**Supplementary Fig. 2B, Supplementary Table 14**).

Given the large phylogenetic distance covered in this study, most orthogroups are represented by multiple genes within a single species. The gene-count per species was between 1 to 8 genes for 96% of the orthogroups for the average species (**Supplementary Fig. 3, Supplementary Table 15**). This could lead to contrasting hypoxic responses within one orthogroup. However, only 3.066% of DEGs-containing orthogroups (|log_2_FC > 1| & Padj. < 0.05) had both up- and down-regulated genes within a single species (**Fig. 1D**, **Supplementary Table 16**). This suggests the need to co-ordinately regulate gene expression among high similarity genes within a species rather than compensatory mechanisms or switches between isoforms. Additionally, it allows orthogroups to act as a vehicle to directly compare transcriptomic responses among distantly related species and identify contrasting and shared responses of the seven species considered in the analysis. We defined an orthogroup as ‘up’ or ‘down’ regulated in a species when at least one DEG (|log_2_FC > 1| & Padj. <0.05) was present. Overall, the transcriptional response to hypoxia was confirmed as strongest, in terms of number of regulated orthogroups, in *O. sativa* and *A. thaliana* (**Fig. 1D**). This analysis revealed many orthogroups (classified as conserved, gained, lost and clade-specific) as downregulated by hypoxia in the two angiosperm species (**Fig. 1D-E**). Also considerable downregulation was observed in *M. polymorpha*. In contrast, downregulation was strongly attenuated in the other species, and virtually absent in *P. patens* and *S. moellendorfii*. Furthermore, the strong regulation in angiosperms, and weaker regulation in other species was not restricted to newly evolved, lost or clade specific-orthogroups, but occurred equally among conserved orthogroups.

### The molecular function of conserved hypoxia-regulated orthogroups

We analysed the extent of overlap of function in regulated orthogroups across species. To this end we compared the similarity of up- and down-regulated orthogroups between all species pair combinations, normalised by the total number of shared orthogroups (**Supplementary Fig. 2B**). As expected, we found the most overlap in regulation in more closely related species, here tracheophytes showed more similar regulation than the bryophytes (**Fig. 2A**). However, this was not the case for repressed orthogroups: in *S. moellendorfii* only few genes were downregulated in response to hypoxia, while in *P. vittata* a unique set of orthogroups, mainly belonging to the ‘gained’ and ‘lost’ categories, was repressed (**Fig. 1E**). Both pteridophyte species, *P. vittata* and *E. hyemale*, shared a similar number of repressed orthogroups with angiosperms. Angiosperms shared between themselves the highest proportion of up- and down-regulated orthogroups.

**Figure 2.**
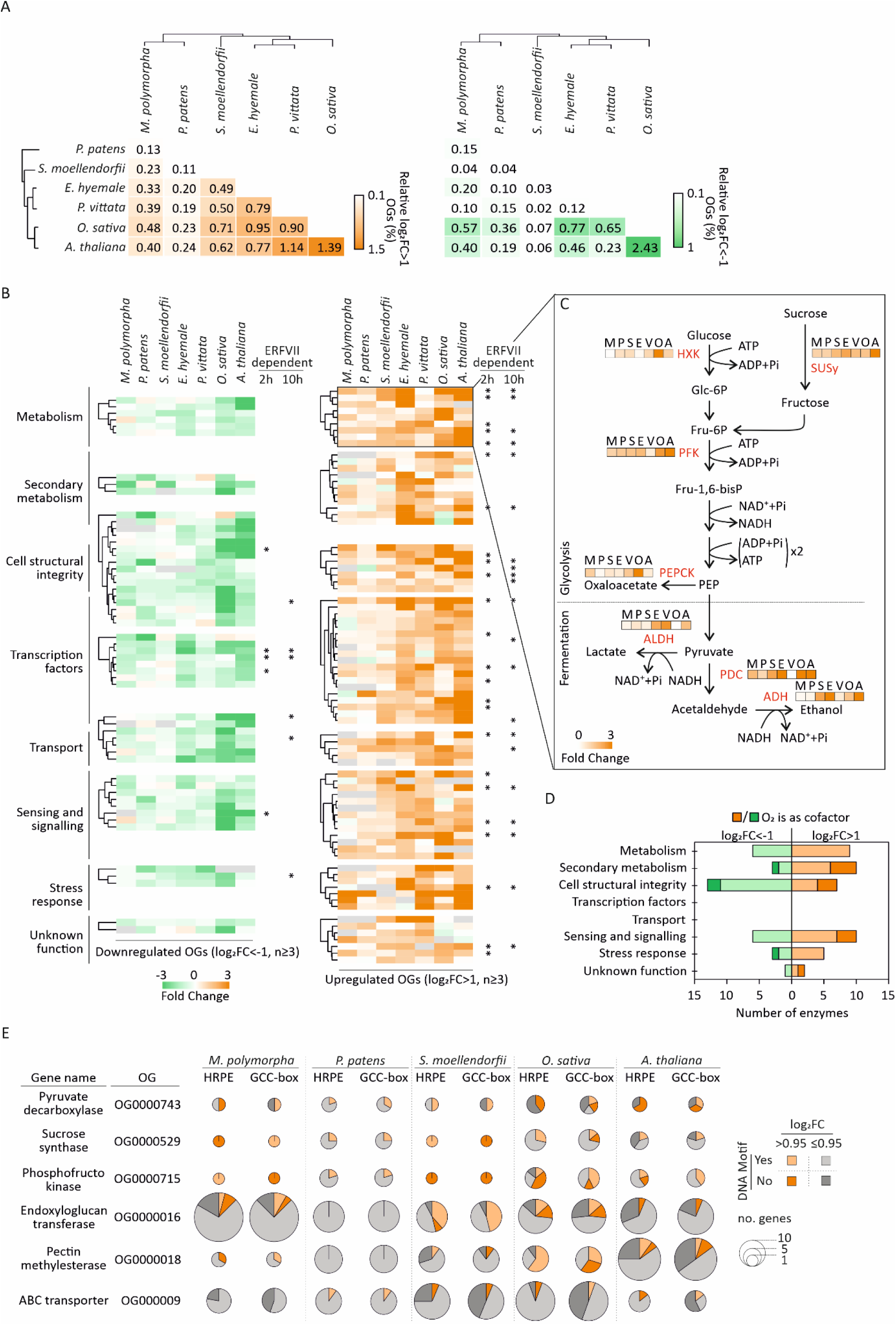
The evolution of the transcriptional response to hypoxia in land plants. **A**) Pairwise comparison of the differentially regulated (|log_2_FC|>1) orthogroups in the six species analysed in this study, normalised by the total number of shared orthogroups for each pair. **B**) Heatmaps representing shared upregulated (log_2_FC>1) and downregulated (log_2_FC<-1) orthogroups between at least three species, grouped according to their molecular function. Orthogroups whose hypoxia regulation (2 and 10 h treatment) is ERFVII-dependent in *A. thaliana* (**Supplementary Fig. S4**) have been labelled with an asterisk. **C**) Simplified scheme of the glycolytic and fermentation pathways in plants, with highlighted in red the enzymes corresponding to the conserved upregulated orthogroups, across species, included in the ‘metabolism’ category. The degree of upregulation for each orthogroup is given by the heatmap next to the corresponding enzyme. Species names are defined as follow: M (*M. polymorpha*), P (*P. patens*), S (*S. moellendorfii*), E (*E. hyemale*), V (*P. vittata*), O (*O. sativa*) and A (*A. thaliana*). **D**) Number of enzymes, identified among the conserved hypoxia-inducible (log_2_FC>1) or repressed (log_2_FC<-1) orthogroups, that use or not use O_2_ as cofactor. **E**) Pie charts representing the frequency of HRPE or GCC-box *cis*-elements found in promoters of upregulated (log_2_FC>0.95) or unchanged (log_2_FC≤0.95) genes belonging to conserved orthogroups under hypoxia. Each pie chart size is adjusted according to the number of genes included in each orthogroup per species.

We further focussed our analysis of the conservation in the transcriptional response to hypoxia in land plants by looking for orthogroups that showed similar regulation. Since conservation across all species was relatively low we identified the orthogroups with regulation in at least three clades, to capture any similarity in regulation across divisions. By applying this criterion, we identified 78 conserved up-regulated orthogroups and 54 down-regulated orthogroups. We grouped them according to the molecular function of the orthologous *A. thaliana* gene, where this information was available (**Fig. 2B**, **Supplementary Tables 17-18**). We observed orthogroups containing genes involved in primary and secondary metabolism, cell structure integrity, small molecule and ion transport, regulation of mRNA transcription, signal transduction and stress responses (**Fig. 2B**). A function could not be attributed to several orthogroups, which we then categorised as ‘unknown’. Several of the conserved up-regulated orthogroups include proteins defined as part of the ‘core anaerobic response’ in *A. thaliana* (Mustroph et al. 2009). Most of the highly conserved induced orthogroups (up-regulated in six out of seven species) belong to the ‘metabolism’ category, including genes involved in the glycolytic and fermentative pathway, such as sucrose synthase (SUS), ATP-dependent phosphofructokinase (PFK) and pyruvate decarboxylase (PDC) (**Fig. 2C**). Remarkably, several genes included in the conserved induced, but not repressed, orthogroups code for signalling and metabolism enzymes that use molecular oxygen as a co-substrate (**Fig. 2D**, **Supplementary Tables 17-18, 19**). Among transcriptional regulators, we found the ERF and MYB families as universally represented, even though the sequence similarity of upregulated genes within these orthogroups was only limited to the DNA binding domain, as opposed to the high similarity throughout the whole sequence observed for metabolic enzymes (**Supplementary File S1-8**).

A transcriptome analysis between wild type *A. thaliana* and the pentuple *erfVII* mutant (with all ERFVII activity removed; *rap2.12 rap2.2 rap2.3 hre1 hre2*; (Abbas et al. 2015)), subjected to 2 and 10 hours hypoxia (**Supplementary Table 20**), revealed that approximately half of the genes induced after 2 hours hypoxia in the wild type were less expressed in the *erfVII* mutant, whereas the absence of ERFVII activity only affected around one fifth of hypoxia-repressed genes. After 10 hours hypoxia, only one fifth of the induced genes exhibited ERFVII-dependency, suggesting the involvement of additional mechanisms at later time points to control transcript abundance (**Supplementary Fig. 4**). Several of the conserved up-regulated orthogroups included *A. thaliana* genes that require the ERFVIIs for induction, in contrast to few of the down-regulated orthogroups that exhibited this feature (**Fig. 2B**, **Supplementary Tables 17-18**) (Giuntoli et al. 2017; Zubrycka et al. 2023).

ERF and MYB orthogroups were the most conserved among hypoxia-induced transcription factors, as they were found up-regulated in at least six out the seven species considered in this study. Therefore we tested for enrichment of their corresponding promoter binding elements among differentially upregulated genes. At this genome-scale resolution we only found significant enrichment of the HRPE motif in the angiosperms and the GCC box in *P. patens* and *S. moellendorfii* (**Supplementary Fig. 5**, **Supplementary Tables 21-25**). However, focussing on the presence of the consensus DNA sequences in the promoters of genes belonging to the orthogroups with similarly conserved regulation revealed a more nuanced picture (**Fig. 2E**, **Supplementary Tables 26-30**). The DNA sequences generically recognised by ERF proteins (GCC-box, (Hao, Ohme-Takagi, and Sarai 1998)) and the ERFVII-specific Hypoxia Responsive Promoter Element (HRPE motif, (Gasch et al. 2016)) were individually present in most, but not all, of the induced-gene promoters of the orthogroups set. They were sometimes found in promoters that are not hypoxia-responsive, indicating that their presence is not sufficient for regulation by the ERFVIIs. Either HRPE or GCC elements were rarely present in *P. patens* promoters, while often found in *M. polymorpha* and the tracheophytes considered in this study. The two DNA elements bound by R2R3-MYB TFs (MYB-I and II, (Kelemen et al. 2015)) were enriched in up-regulated orthogroups of *O. sativa* and *A. thaliana* promoters, and more wide-spread in down-regulated orthogroups promoters, although absent in *P. patens* (**Supplementary Fig. 6A**). The DNA element recognised by membrane-tethered NAC transcription factors (MDM, (De Clercq et al. 2013), which have been described to attenuate the oxidative stress associated with submergence and reoxygenation in *A. thaliana* (Giuntoli et al. 2017), was not enriched in our cross-species dataset (**Supplementary Fig. 6B**).

### ERFVII evolution in land plants

The enrichment of ERF-recognised DNA elements and the conservation of ERF-dependent genes among hypoxia-inducible orthogroups across land plants prompted us to investigate how widespread involvement of ERFVIIs is in the regulation of the transcriptional response to acute hypoxia in the green lineage. Since our orthogroup analysis could not fully distinguish groups within the ERF subfamily (Nakano et al. 2006), we analysed best reciprocal hits using the *A. thaliana* RAP2.12 protein as a bait against the genome predicted proteomes of 104 land plant species (12 bryophytes and 92 tracheophytes) and two algae species (one chlorophyte and one charophyte). For each tracheophyte species considered, we could retrieve at least one protein sharing similarity with *A. thaliana* ERFVIIs, that extended outside of the AP2 domain (**Supplementary Table 31**). Conversely, for only a few bryophyte species our analysis could identify sequences similar to *A. thaliana* ERFVIIs, while for the majority of bryophytes and algae the sequence similarity was limited to the AP2 DNA binding domain (**Supplementary File S9**, **Supplementary Tables 32-33**). We used all the tracheophyte ERFVII-like proteins retrieved in this study to generate a phylogenetic tree, where sequence similarity recapitulated species subdivision in four clades: lycophytes, ferns, gymnosperms and angiosperms (**Fig. 3A**). The sequences in the fern branch of our tree could not separate eusporangiate and leptosporangiate species, suggesting that ERFVII divergence predated the separation of the two clades (**Fig. 3A**). Additionally, a group of ERFVIIs in the *Equisetum* genus shared such a high sequence similarity with gymnosperm sequences to be included in their same branch of the tree (**Fig. 3A**, **Supplementary File S10**). An in-depth inspection of the ERFVII-like protein N-terminus (first 15 aa) confirmed the extreme sequence conservation in spermatophytes reported before (Holdsworth and Gibbs 2020). In particular, Cys2, required for oxygen-regulated degradation (Zubrycka et al. 2023) is present in lycophyte and fern ERFVII-like proteins, but the N-terminal sequence diverges in subsequent positions, exhibiting high variability from the 5^th^-6^th^ residues onwards (**Fig. 3B**). Gly is the preferred residue in the 4^th^ position across all tracheophytes, while Arg predominates in the 3^rd^ position in lycophytes and was often also found in eusporangiate ferns (**Fig. 3B**). Gly substitutes the Arg3 residue in ferns and spermatophytes, generating the MCGGAI consensus previously reported for angiosperms. Remarkably, leptosporangiate ERFVII-like proteins often contain a Trp in the 3^rd^ position (**Fig. 3B**). We tested whether the ERFVII-like proteins from Lycophytes and ferns are substrates of the PRT6 N-degron and Ubiquitin Proteasome (UPS) pathways (**Fig. 3C**). We cloned the coding sequence of ERFVII-like protein genes from *S. moellendorfii, E. hyemale* and *P. vittata* downstream of a dihydrofolate reductase (DFHR) and Ubiquitin (UBQ) fusion expressed under the control of the CaMV 35S promoter (Varshavsky 2011; Zubrycka et al. 2023). This construct allows testing of the stability of ERFVII proteins after they are co-translationally released from the precursor polypeptide by the cleavage activity of constitutive deubiquitinases (**Fig. 3C**). A pulse-chase experiment using coupled transcription/translation in rabbit reticulocyte extracts showed that these proteins were unstable, but that the half-life was increased in the presence of the proteasome inhibitor Bortezomib (BZ) or in Cys2Ala constructs (**Supplementary Fig. S7A-B**), indicating that their stability is regulated by Cys2. PRT6 and UPS regulated degradation was assessed *in vivo* in *A. thaliana*: when transformed in the *A. thaliana erfVII* mutant, the proteins were highly unstable and failed to accumulate in seedling tissues. However, accumulation of all ERFVII-like proteins was increased either by genetic inactivation of the N-recognin PRT6 (in the *erfVII prt6* sextuple mutant) or through application of the proteasome inhibitor BZ (**Fig. 3D**). Similarly, scavenging of NO with cPTIO or low oxygen conditions (submergence or hypoxia) both led to stabilisation of the proteins (**Fig. 3D**, **Supplementary Fig. S7C-D**). Together, these results confirm that lycophyte and fern ERFVII-like proteins are substrates of the O_2_ and NO-dependent PCO N-degron pathway in *A. thaliana*. Given the conserved nature of this pathway in eukaryotes, we suggest that these act as bona fide PCO N-degron pathway substrates also in their species of origin.

**Figure 3.**
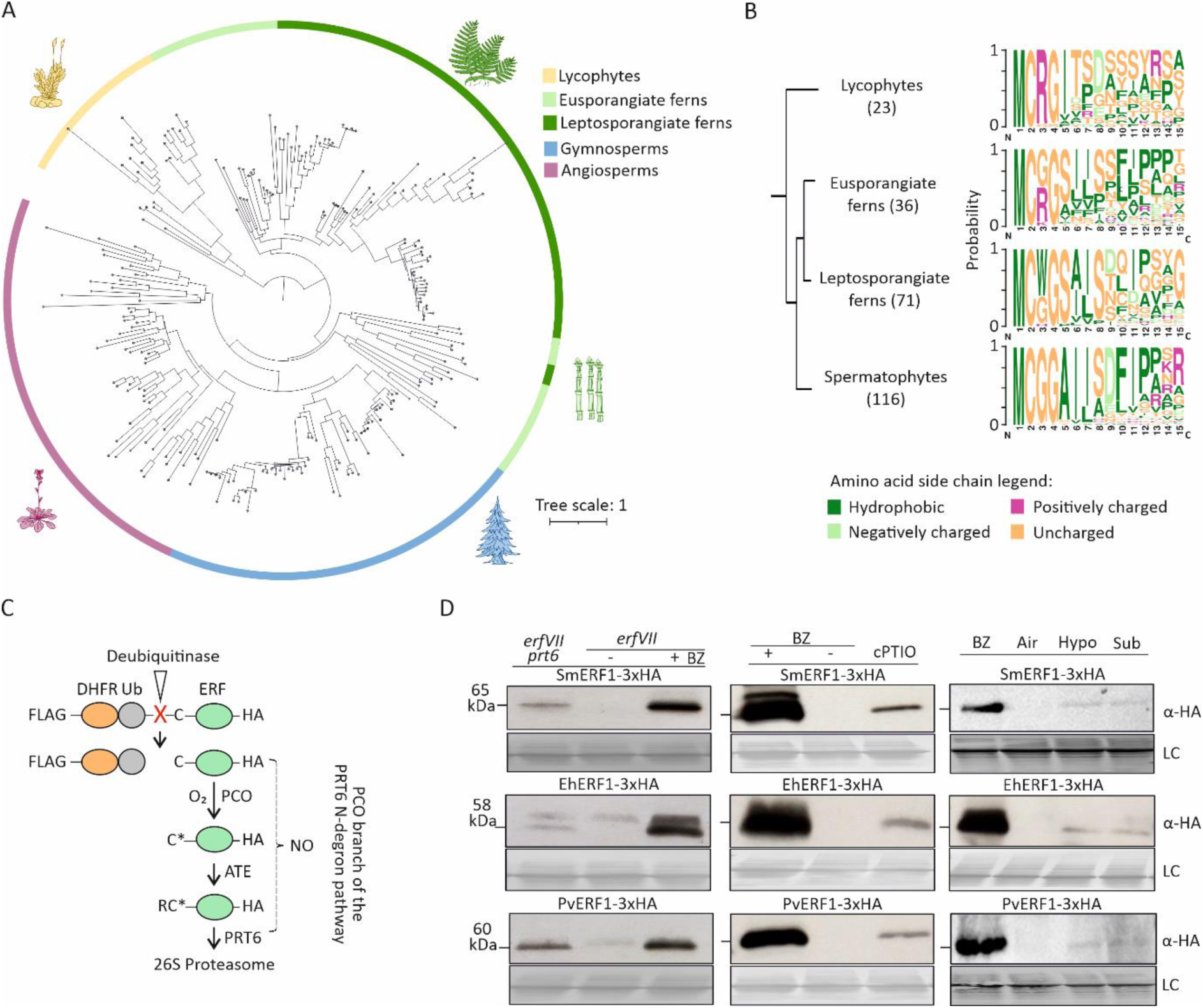
Evolution of the Group VII of Ethylene Responsive transcription Factors (ERFs) in tracheophytes. **A**) Rooted phylogenetic tree of ERFVII orthologues in tracheophytes (lycophytes, eusporangiate ferns, leptosporangiate ferns, gymnosperms and angiosperms). **B**) Motif logo of the first 15 N-terminal amino acids of the ERFVII sequences used to construct the phylogenetic tree shown in **A**). The number of species included in each group is highlighted in brackets. **C**) Schematic representation of the ubiquitin fusion technique construct (Zubrycka et al. 2023) used to study the protein stability of different ERFVIIs, tagged with 3xHA for immunoblot detection, through the PCO branch of the PRT6 N-degron pathway. **D**) Western blot analysis of SmERF1^3xHA^, EhERF1^3xHA^ and PvERF1^3xHA^ abundance in control (DMSO), bortezomib (BZ), NO-scavenger (cPTIO), hypoxia (2h, 0.1% v/v O_2_) or submergence (1h) treated *erfVII* or *erfVII prt6* mutant seedlings. Loading control (LC) shown corresponds to a Ponceau staining of total loaded proteins on Western blots.

### Non-ERFVII ERF proteins have been recruited as substrates of the PCO N-degron pathway in bryophytes

Assessed algal genomes did not contain any ERFVII-like proteins, or ERF proteins with Cys2 (**Supplementary Table 33**). Within the Marchantiophyta, *M. polymorpha* (MpERF18, Mapoly0092s0058), together with other Marchantiopsida and Jungermannopsida species include proteins with similarity to angiosperm ERFVIIs, although none contains Cys2 (**Supplementary Fig. S8, Supplementary Table 34**). While these sequences clustered with *A. thaliana* and *S. moellendorfii* ERFVIIs, similar proteins were absent in moss (*P. patens*) and hornwort (*Anthoceros punctatus*) (**Supplementary Fig. S9**). However, we found a number of ERF proteins with conserved Cys2 (Cys2-ERFs) (**Supplementary Table 35**). A genome-wide analysis of the whole ERF family in *M. polymorpha, P. patens* and *S. moellendorffii,* and including the hornwort *Anthoceros punctatus,* revealed a conserved set of Cys2-ERFs with strong sequence similarity to ERF Group X, Group IV (*M. polymorpha*) and Group VIII (*P. patens*) (**Fig. 4A**). The conserved Group X Cys2-ERFs exhibited an extended N-terminal consensus where positively charged residues preferentially occupied the third position, a feature shared by ERFVIIs in lycophytes and eusporangiate ferns (**Fig. 3B**, **4B**). We tested whether *M. polymorpha* and *P. patens* Cys2-ERFs are substrates of the PCO N-degron pathway. The combination of *in vitro* and *in vivo* assays also used for tracheophyte ERFVII-like proteins (**Fig. 3C**) indicated that these Cys2-ERF proteins are unstable unless the proteasome is inhibited or N-degron processing is genetically (*erfVII prt6* background) or chemically prevented through NO scavenging with cPTIO (**Fig. 4C, Supplementary Fig. S10**). However, different from tracheophyte sequences, we could not observe protein stabilisation in response to hypoxia or submergence, whereas an *A. thaliana* ERFVII RAP2.3-based reporter (Zubrycka et al. 2023) exhibited the expected stabilisation in both conditions (**Supplementary Fig. S7C-D**). This suggests that the apparent insensitivity of bryophyte Cys2-ERFs to hypoxia could be due to sustained Cys oxidation by *A. thaliana* PCOs under oxygen limiting conditions. Therefore, we speculated that, while bryophyte Cys2-ERFs proteolysis is insensitive to hypoxia, affinity of tracheophytes ERFVIIs for PCOs is optimally reduced through Gly and positively charged residues to disfavour oxidation, and subsequent degradation, under hypoxia. Indeed, a fusion between a 50 aa-long N-terminal fragment of *A. thaliana* RAP2.12 and the fluorescent protein Citrine exhibited increased nuclear accumulation in hypoxia when expressed in *M. polymorpha* (**Fig. 4D**, **Supplementary Fig. S11**), confirming the intrinsic property of tracheophyte ERFVIIs to be regulated by oxygen availability. We additionally confirmed the independence of the *M. polymorpha* Cys2-ERF (MpERF23, Mapoly0293s0001) from oxygen at the functional level: CRISPR-mediated inactivation of this gene did not affect the expression of hypoxia responsive genes in *M. polymorpha* (**Fig. 4E**).

**Figure 4.**
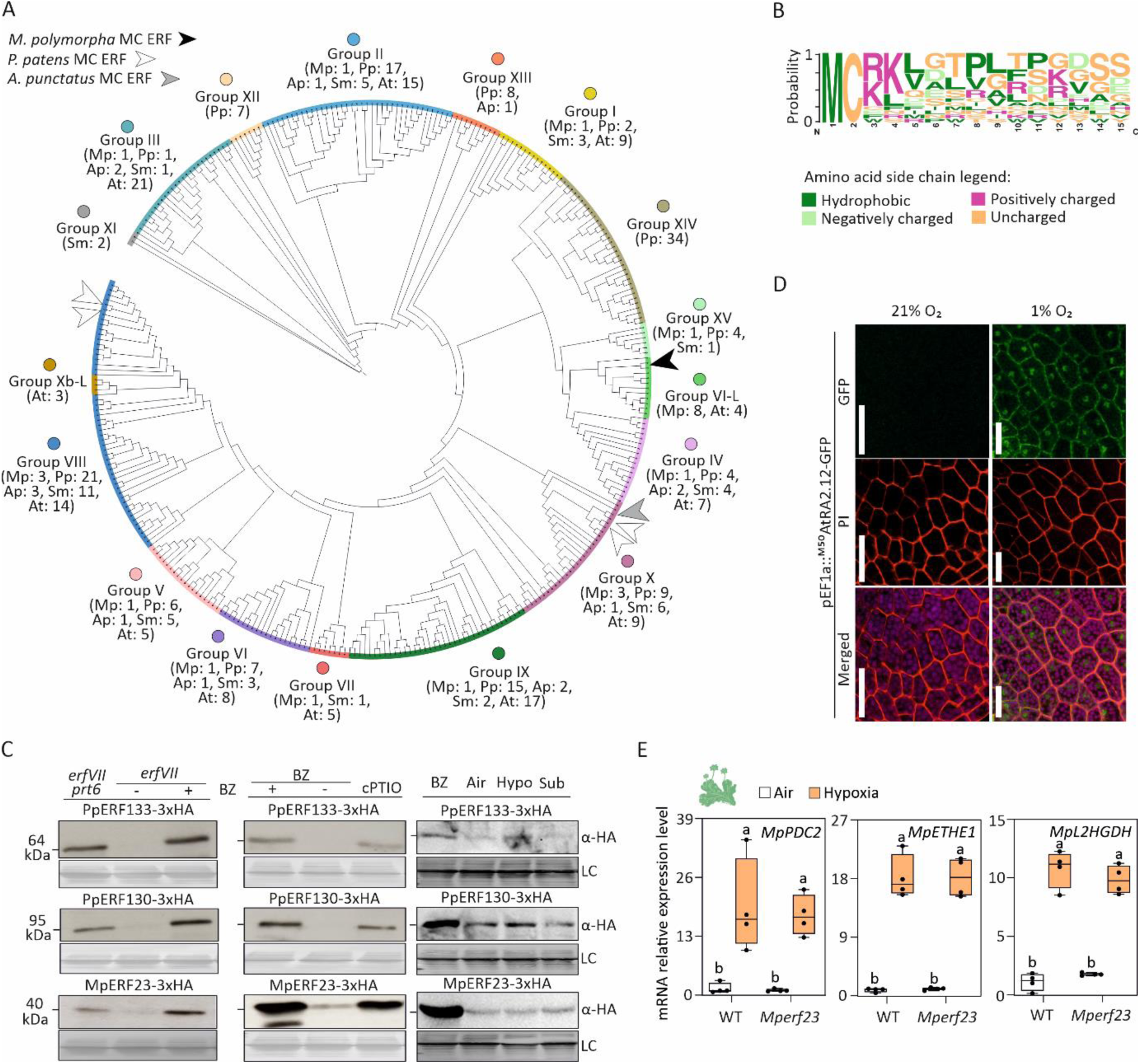
Characterisation of Cys2-ERF proteins in bryophytes. **A**) Unrooted phylogenetic tree of ERF proteins from bryophytes (*A. punctatus*, *M. polymorpha* and *P. patens*), lycophyte (*S. moellendorfii*) and angiosperm (*A. thaliana*). ERFs classification is based on (Nakano et al. 2006). White, grey and black arrow heads represent Cys2 ERFs in *P. patens*, *A. punctatus* and *M. polymorpha*, respectively. **B**) Motif logo of the first 15 N-terminal amino acids of the Cys2-ERF Group X sequences retrieved in bryophytes (liverworts, mosses and hornworts). **C**) Western blot analysis of PpERF133^3xHA^, PpERF130^3xHA^ and MpERF23^3xHA^ abundance in control (DMSO), bortezomib (BZ), NO-scavenger (cPTIO), hypoxia (2h, 0.1% v/v O_2_) or submergence (1h) treated *erfVII* or *erfVII prt6* mutant seedlings. Loading control (LC) shown corresponds to a Ponceau staining of total loaded proteins on Western blots. **D**) Confocal images of the subcellular localization of ^1-50^AtRAP2.12-Cytrine, overexpressed in *M. polymorpha* 13-day-old thalli in air (21% ambient O_2_) and hypoxia (1% v/v O_2_) (**Supplementary Fig. S11**). White arrowheads indicate the position of nuclei in the GFP channel. **E**) Effect of MpERF23 on *MpPDC2*, *MpETHE1* and *MpL2HGDH* expression levels in air and hypoxia (state level) in wild-type compared to *Mperf23* knockout 13-day-old *M. polymorpha* thalli. Data are presented as means ± SD (n = 4). Different letters indicate statistically different averages as assessed by two-way ANOVA followed by the Holm-Sidak posthoc test (P < 0.05).

### Evolution of Cys2-ERFs as oxygen-dependent transcriptional regulators

We set out to test the ability of the bryophyte and tracheophyte Cys2-ERFs, which are substrates of the PCO N-degron pathway, to substitute endogenous *A. thaliana* ERFVIIs in activating hypoxia-related gene expression in seedlings. Under submergence no significant changes in expression of conserved hypoxia-activated genes were observed in plants containing Cys2-ERF transgenes compared to the *erfVII* mutant (**Fig. 5A**). However, genetic inactivation of the N-degron pathway via *prt6* in the *erfVII* background (*erfVII prt6*) led to increased expression of *PDC1* and *ADH1* (**Fig. 5B**). We speculated that absence of complementation of the *erfVII* response to submergence by non-spermatophyte Cys2-ERFs may be due to an inability of these proteins to interact with the transcriptional machinery in *A. thaliana*. To test this hypothesis, we fused the *A. thaliana* RAP2.12 activation domain (aa 340-358, (Bui et al. 2015)) to each Cys2-ERF and measured the ability to transactivate an HRPE or GCC-box containing synthetic promoter in *A. thaliana* mesophyll protoplasts (**Fig. 5C**). The *P. patens* Cys2-ERFs, *S. moellendorffii* and *P. vittata* ERFVIIs could significantly induce the luciferase reporter gene when driven by the HRPE at a reduced level compared to RAP2.12, but not the GCC-box containing promoter (**Fig. 5D**). This suggested that HRPE-binding is not limited to ERFVIIs and that other phylogenetically distant Cys2-ERFs possess specificity for this DNA motif (**Fig. 2D**). To test the ancestral role of HRPE in mediating hypoxia responsiveness *in cis*, we cloned three *S. moellendorffii* genomic regions upstream of hypoxia induced genes (*ADH_prom_*, *PDC_prom_* and *HB_prom_*) and tested whether they could be activated by RAP2.12 or SmERF1 in *A. thaliana* protoplasts. RAP2.12 activated all three promoters, while SmERF1 could activate only two of them, leading us to hypothesize that regulation of hypoxia responsive genes in lycophytes is likely mediated by ERFVIIs, as demonstrated for angiosperms (Licausi et al. 2011), but this apparently did not require the presence of an HRPE or GCC-box motif (**Fig. 5E-G**). Although our assay could not test for direct DNA-binding, this result suggested that the role of HRPEs is contingent to the DNA context in which they occur. A protoplast assay also showed that the *M. polymorpha* ERFVII-like protein that is similar to angiosperm ERFVIIs, but does not contain Cys2, can regulate expression through the synthetic HRPE promoter, but not the GCC-box (**Supplementary Fig. S12**), corroborating the hypothesis of an early partnership between ERFVIIs and HRPE.

**Figure 5.**
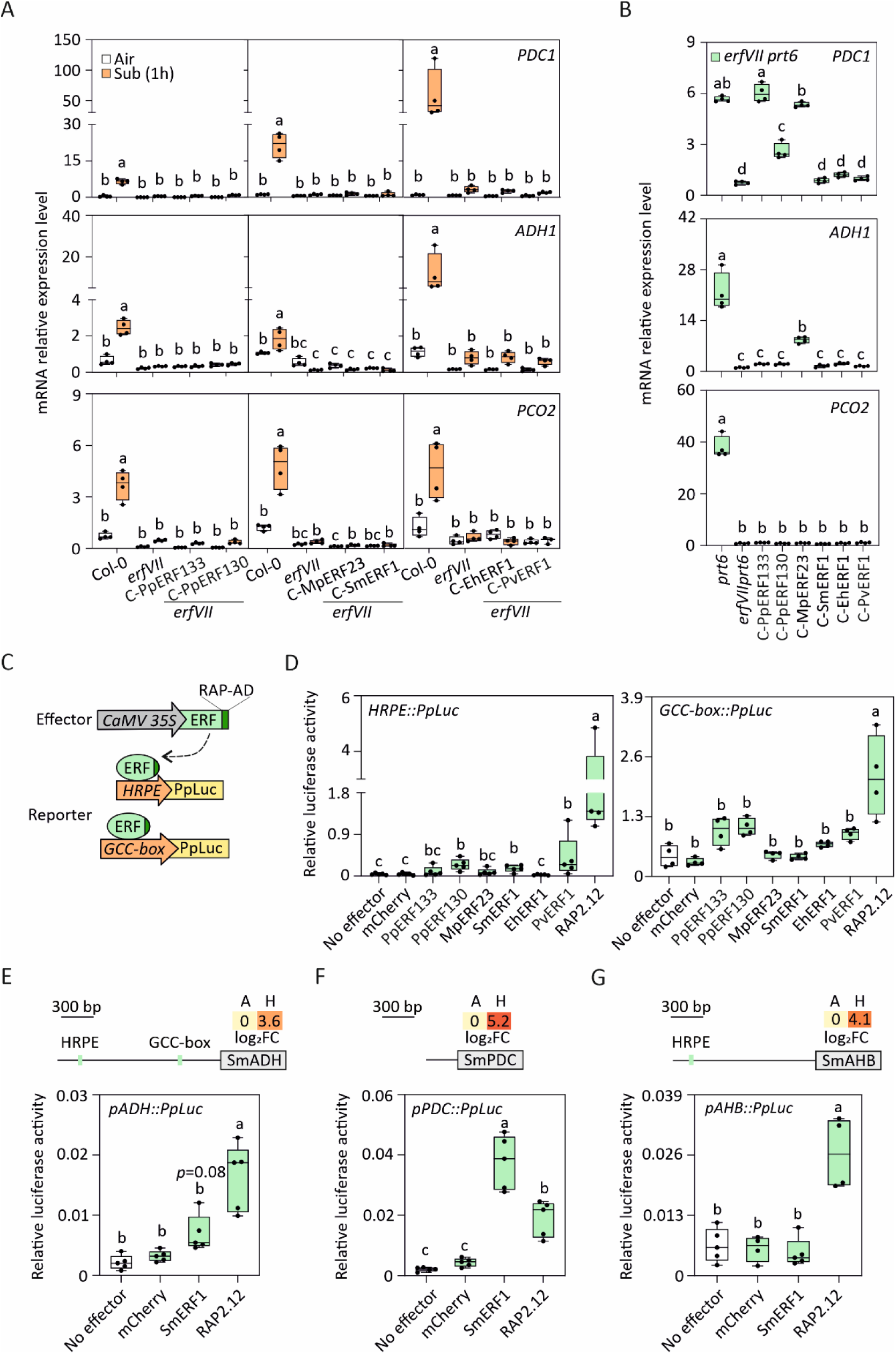
Potential of Cys2-ERFs and non-angiosperm ERFVIIs to regulate hypoxia-responsive genes in *A. thaliana*. **A**) Effect of Cys2-ERFs or ERFVIIs on the expression of *PDC1*, *ADH1* and *PCO2* in 7-day-old *A. thaliana erfVII* seedlings following one hour of dark submergence compared to aerobic conditions. **B**) Effect of Cys2-ERFs or ERFVIIs on the expression of *PDC1*, *ADH1* and *PCO2* in 7-day-old *A. thaliana erfVII prt6* seedlings. **C**) Scheme representing the effector (ERF coding sequence fused to the AtRAP2.12 activation domain, under the control of the constitutive CaMV 35S promoter) and reporter (firefly luciferase coding sequence under the control of the synthetic, hypoxia-inducible promoter HRPE or ERF-specific GCC-box) constructs used in the protoplasts transactivation assay in **D**. **D**) Effect of Cys2-ERFs or ERFVIIs on the synthetic promoters HRPE and GCC-box measured as relative firefly luciferase activity in *A. thaliana erfVII prt6* mesophyll protoplasts. Data are presented as means ± SD (n = 5). Different letters indicate statistically different averages as assessed by one-way or two-way ANOVA followed by the Tukey’s method or the Holm-Sidak posthoc tests, respectively (*p*-value<0.05).

## Discussion

This study explored the conservation and diversification of the transcriptional response to hypoxia in land plants and the evolution of the ERFVII transcription factors, which play a major role as transducers of hypoxic signalling in angiosperms. While the enzymatic components of the PCO N-degron pathway are ubiquitously present in the plant kingdom, inherited from the last common ancestor of eukaryotes (Holdsworth and Gibbs 2020; Weits et al. 2023), we found that the ERFVII substrate transducers are specific to tracheophytes, where they orchestrate gene expression changes in low oxygen. This places the evolution of the ERFVIIs at least 250 million years later than the metazoan Hypoxia Inducible Factor (Mills et al. 2018; Erwin et al. 2011). Recruitment of different ERF family members as N-degron pathway substrates occurred independently several times during land plant evolution, as observed in liverworts and mosses; however, it is unclear whether these TFs contribute significantly to mount a transcriptional response to hypoxia (**Fig. 6**). ABR1, a highly conserved Cys2-ERFX, was shown to escape N-degron proteolysis in *A. thaliana*, where it regulates stress signalling and wound repair mechanisms (Baumler et al. 2019).

**Figure 6.**
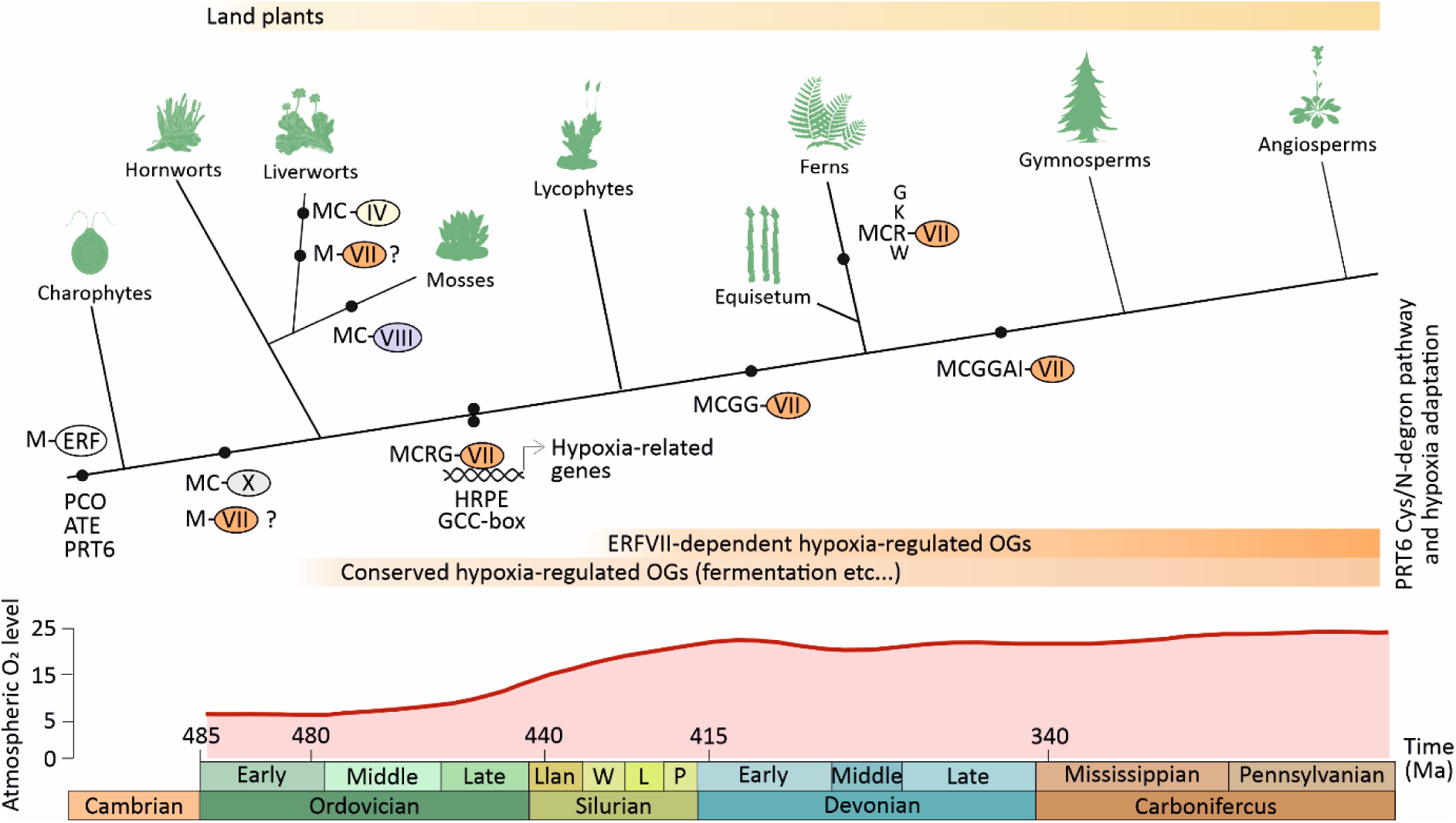
Evolution of Cys2-ERFs as transducers of hypoxia signalling in land plant. The key enzymatic components of the PCO N-degron pathway (PCO, ATE and PRT6) likely evolved in the last common ancestor of Eukaryotes. Cys2-ERFs arose multiple times in land plants and ERFVIIs evolved O_2_-dependent degradability in vascular plants. The Cys2-ERF proteins in bryophytes belong to Group X, VII (mosses) and IV (liverworts). In ferns, the extended N-terminal sequence of ERFVIIs (CMVII motif) shows variability compared to spermatophytes and lycophytes. ERFVII-like proteins are found in liverworts, although these proteins do not carry a Cys in second position. The recruitment of the cis-elements HRPE and GCC-box in promoters of hypoxia-regulated genes is a feature of tracheophytes.

The orthogroups whose expression was altered by reduced oxygen availability varies across species, as expected due to anatomical and ecological divergence; nevertheless, a small set of induced orthogroups was conserved in all plant species investigated in this study (**Fig. 2B**). This ‘core hypoxia-response orthogroups’ included genes involved in metabolic reconfiguration towards glycolysis and fermentation, cell wall modifications and molecular transport that were up-regulated similarly (**Fig. 2B**). Indeed, ethanol production was previously identified as a highly conserved adaptation to low oxygen in plants (Bui et al. 2019). Moreover, many of the induced metabolic genes code for enzymes that use oxygen as a co-substrate and therefore their induction may be interpreted as a compensation mechanism, a phenomenon already reported for the oxygen sensor PCO (Weits et al. 2023). Our analysis also identified a set of hypoxia-regulated orthogroups for which a function could not be attributed due to lack of similarity to known proteins. They should be studied to fully understand the mechanisms of adaptation to hypoxia in plants.

Our findings suggest the potential existence of an ancestral hypoxia signalling mechanism common to all land plants. Interestingly *A. thaliana* is still able to activate a transcriptional response in the absence of ERFVIIs (**Supplementary Figure S4**). Evidence for conservation in signalling events is also suggested by the induction of MYB and ERF orthologues in most of the species we investigated. The non-flowering species exhibited a reduced transcriptional response to hypoxia compared to angiosperms. This can be interpreted as a demonstration of a less pronounced sensitivity towards this stress or, alternatively, could suggest the existence of non-transcription associated regulation. The stronger transcriptional response to hypoxia observed in *A. thaliana* and *O. sativa* (**Fig. 1 and 2A**) suggests an expansion of the sphere of influence of ERFVIIs over plant cell physiology through the recruitment of newly evolved and conserved orthogroups. Our phylogenetic analysis revealed a trend for ERFs to be recruited as substrates of the PCO-branch of the PRT6 N-degron pathway with the independent evolution of Cys2-ERFs from different groups within the family multiple times in different plant divisions. The conservation of Cys2 in ERFs outside angiosperms suggests that these are important for the response to conditions that require PRT6 N-degron pathway regulated proteolysis. When heterologously expressed in *A. thaliana*, non-vascular Cys2-ERFs were stabilised by NO scavenging but not hypoxia or submergence (**Fig. 4C**). This confirms the previously reported independence of PCO-mediated cysteine oxidation from cellular NO levels, and indicates a requirement of NO for steps in the PRT6 N-degron pathway downstream of PCO (Puerta et al. 2019; Gibbs and Holdsworth 2020). On the other hand, it suggests a potential role for Cys2-ERFs from non-vascular clades specifically in regulating environmental or developmental conditions that decrease NO availability, as shown in flowering plants (Vicente et al. 2017b; Vicente et al. 2019). For example NO synthesis via nitrate reductase is reduced under drought and high salinity stress and shoot apical meristem establishment is responsive to lowered NO levels (Gibbs et al. 2014; Vicente et al. 2017a; Zeng et al. 2023).

We observed that ERFVIIs are only found as Cys2-proteins (and thus substrates of the PCO N-degron pathway) in vascular plants (**Fig. 6**, (Weits et al. 2023)). Their N-terminal consensus sequence (CMVII motif), which does not exist in non-vascular plant ERFs, exhibits strong sequence conservation beyond Cys2 (**Fig. 3B**). This may represent an optimised sequence for interaction with N-degron enzymes, or other components of the hypoxia response transcriptional machinery. Importantly, ERFVIIs presence only in vascular plants correlates with the recruitment of genes to constitute a core transcriptional response previously identified in angiosperms (**Supplementary Tables 1-5**) (Mustroph et al. 2010). When tested in *A. thaliana* as a heterologous host, non-spermatophyte ERFVIIs could not restore fast induction of low oxygen responsive genes (**Fig. 5A**), which we attribute to the absence of complementation to division-specific components of the transcription machinery. Indeed, fusion of the angiosperm activation domain of RAP2.12 (Bui et al. 2015) to Cys2-ERFs from non-vascular clades led to significant induction of a hypoxia-responsive reporter in protoplasts (**Fig. 5D**). Moreover, transient transformation of *A. thaliana* protoplasts with the *S. moellendorfii* ERFVII and hypoxia-responsive promoters indicated conservation of both *cis* and *trans*-regulatory components (**Fig. 5E**). We therefore conclude that ERFVIIs likely acquired their role as transducers of the hypoxia response in the last common ancestor of vascular plants. The presence of similar ERFs in liverworts (that do not contain Cys2 or the CMVII motif) suggests that they might have originated from an older ancestor in non-vascular plants which was lost in sister lineages (**Supplementary Table 34**, **Fig. 6**). However, the presence of several unique motifs in ERFVIIs also indicates a de-novo origin at the junction between vascular and non-vascular plants.

Although non-vascular plants do not have hypoxia-stabilised ERFVIIs, they do adjust gene expression to hypoxia. A direct oxygen sensing mechanism could be responsible for the transcriptional response in bryophytes. This could involve a non-ERF N-terminal Cys-transcriptional regulator, converted to a substrate of the PCO N-degron pathway by endoproteolysis. An alternative biochemical strategy for direct oxygen sensing in bryophytes could rely on 2-oxoglutarate dependent dioxygenases, such as those identified in the animal kingdom (Ivan et al. 2001; Chakraborty et al. 2019; Batie et al. 2019). The induction of O_2_-consuming enzymes in *M. polymorpha* and *P. patens* could hint at the existence of a feedback mechanism where the O_2_-sensing enzymes are hypoxia inducible as in the case of PCOs and PHDs (Weits et al. 2014; D’Angelo et al. 2003). Less direct O_2_-sensing mechanisms could be responsible for gene induction upon hypoxia. These could involve variations in reactive oxygen species levels, cytosolic Ca^2+^ concentrations and changes in lipid saturation that contribute to low O_2_ signalling in *A. thaliana* (Schmidt et al. 2018; Pucciariello and Perata 2017; Fan et al. 2023).

The ERFVIIs are the most ancient PCO N-degron substrates identified to date. They are present as ubiquitously highly conserved in all vascular plants, while VERNALIZATION 2 (VRN2) and LITTLE ZIPPER 2 (ZPR2) are angiosperm-specific, suggesting a derived role specific to angiosperms (Gibbs et al. 2018; Labandera et al. 2021). It remains puzzling why ERFVIIs evolved in vascular plants. A recent report (Zubrycka et al. 2023) demonstrated that in *A. thaliana* they are essential to maintain an extended root system in waterlogged soils, and contribute to root growth under normal conditions. This indicates a strong requirement for ERFVII function for root development and survival in soil. Complex soils first appeared in the terrestrial environment at the time of the first appearance of plants with true roots (Spencer et al. 2022). An important property of soils is the capacity to provide changes in water saturation levels. We speculate that the ERFVIIs might have arisen to ensure survival of the root, a developmental innovation evolutionarily associated with plant vascularisation (Hetherington and Dolan 2018). This feature might have been essential for soil exploration under changing water saturation levels, and therefore contributed to successful habitat colonisation.

## Online Methods

### Plant material and growth conditions

*Marchantia polymorpha* thalli or gemmae, assession Cambridge-2 (Cam-2), *Selaginella moellendorfii* and *Pteris vittata* shoots were cultivated *in vitro* conditions on half-strength MS medium (Murashige and Skoog 1962), supplemented with 0.5% (w/v) sucrose and 0.8% (w/v) plant agar. *Physcomitrium patens* (kindly provided by Prof. Tomas Morosinotto, University of Padua) was grown on Knop medium (Reski and Abel 1985). *M. polymorpha* thalli were sub-cultured from single gemmae while *P. patens*, *S. moellendorfii* and *P. vittata* plantlets from shoot cuttings. Plants were grown under controlled conditions: 50–60 µmol m−2 s−1 white light, 16:8h day:night photoperiod, at 22 °C. *Equisetum hyemale* plants were grown in soil (1:3 ratio of peat-free compost soil and vermiculite), under controlled environmental conditions with a photoperiod of 16:8h, temperature of 20:18°C light:dark and white light supplemented with 10 μm far red light.

The *A. thaliana erfVII* and *erfVII prt6* mutants were described previously (Abbas et al. 2015). The Col-0 accession was used as a wild-type reference. Transgenic Col-0 *A. thaliana* seeds carrying pUFT:RAP2.3^3HA^ were described previously (Zubrycka et al. 2023). *A. thaliana* seeds were sown in moist peat-free compost and vermiculite (1:3) soil, stratified at 4 °C in the dark for 48 h and germinated at 22:18 °C day:night under 16:8h photoperiod. For *in vitro* seedlings cultivation, seeds were surface sterilized and, subsequently, sown in square plates on half-strength MS medium supplemented with 1% (w/v) sucrose and 0.8% (w/v) plant agar. After cold treatment, plates were positioned vertically and seeds germinated under 16:8h day:night photoperiod, at 22 °C. *M. polymorpha* transgenic line pEF1a:^M50^AtRAP2.12-GFP was available in the laboratory.

### Generation of various vectors

The CDS, starting at Cys^2^ or C^2^A, of MpERF23 (Mapoly0293s0001), PpERF133 (Pp3c7_1240V3.1), PpERF130 (Pp3c6_28370V3.1), SmERF1 (SELMODRAFT_73139), EhERF1 (scaffold-JVSZ-2009360) and PvERF1 were cloned from *M. polymorpha*, *P. patens*, *S. moellendorfii*, *E. hyemale* and *P. vittata*, respectively, using Phusion High Fidelity DNA-Polymerase (New England Biolabs). The amplified CDS sequences were cloned through *SacII* digestion into the pCD1 vector (carrying the ^FLAG^DHFR-UBIQUITIN-3xHA cassette, (Zubrycka et al. 2023)) to generate an entry vector. Each entry vector was then recombined into plant transformation gateway vector pH7WG2, using the LR reaction mix II (Life Technologies), to generate an expression vector, as described in (Zubrycka et al. 2023). The *p35S::ccdB-RAP2.12-AD* vector was generated by combining the 35S Cauliflower mosaic virus promoter, a synthetised minimal ccdB cassette, the activation domain of RAP2.12 (RAP2.12-AD, corresponding to 1021-1077 bp (351-358 aa)) and the 35S terminator into the pDGB3alpha2 backbone vector, using the GoldenBraid cloning system (Sarrion-Perdigones et al. 2011). The full CDS of MpERF23, PpERF133, PpERF130, SmERF1, EhERF1, PvERF1 and RAP2.3 were cloned into the gateway entry vector pDONR207 and then subcloned into the destination vector p35S::RAP2.12-AD, through the LR reaction. Primer sequences used to generate these constructs are listed in **Supplementary Table 36**.

### Generation of transgenic plants

The DNA constructs described above were introduced into *erfVII* and *erfVII prt6 A. thaliana* mutants via *Agrobacterium tumefaciens*-mediated transformation using the floral dip method (Clough and Bent 1998). Transgenic seeds were selected on agarised MS medium supplemented with hygromycin B (cat. No. 10687010, Thermo Fischer Scientific). Homozygous plants carrying one single transfer DNA insertion were identified on segregation basis.

### Identification of Cys2-ERFs and ERFVIIs

Identification of ERFVII transcription factor sequences in tracheophytes (lycophytes, ferns and gymnosperms) was performed by searching the ONEKP project database (https://db.cngb.org/onekp/) using the BLAST (tblastn) algorithm (Altschul et al. 1990) and *A. thaliana* RAP2.12 (AT1G53910) protein sequence as query. The retrieved proteins were then align back against *A. thaliana* protein database (www.phytozome.net) to ensure that they represent the closest orthologs. The best *A. thaliana* ERFVII orthologous sequences from angiosperms, together with the five Arabidopsis ERFVIIs (AT1G53910, AT1G72360, AT2G47520, AT3G14230 and AT3G16770) were selected among those identified by (Weits et al. 2023). Orthologues ERFVII sequences from lycophytes, ferns, gymnosperms and angiosperms are listed in **Supplementary Table 31**. Identification of ERF transcription factor sequences, featuring a Cys2 and an AP2 DNA binding domain, in bryophytes and algae was performed as described above for tracheophyte sequences. ERF transcription factors from *C. reinhardtii*, *K. flaccidum*, *M. polymorpha*, *P. patens*, *S. moellendorfii* and *A. thaliana* were retried from the PlantTFDB database (https://planttfdb.gao-lab.org/index.php) (Jin et al. 2017). *Anthoceros punctatus* ERF transcription factors were identified by BLAST search using the orthologous ERF sequences from the hornwort *Anthoceros agrestis* (Li et al. 2020). The complete lists of the above-mentioned ERF sequences are found in **Supplementary Tables 32-33**.

### Phylogenetic analysis

The retrieved ERFVII full sequences from tracheophytes were aligned using the MUSCLE algorithm (Edgar 2004) and used for phylogenetic analysis. After aligning the ERF transcription factor sequences using the MUSCLE algorithm, only the aligned region corresponding to the AP2 binding domain was exported and used for the subsequent phylogenetic analysis. Phylogenetic analysis were performed using the IQ-TREE online application (Trifinopoulos et al. 2016). Phylogenetic trees were visualised, annotated and exported using the Interactive Tree Of Life (https://itol.embl.de) online tool (Letunic and Bork 2021).

### Low oxygen, submergence and chemical treatments

For hypoxia treatments on plantlets, 13-day-old *M. polymorpha*, *P. patens*, *S. moellendorfii*, *P. vittata* and *E. hyemale* plantlets were exposed to hypoxia at 1% v/v O_2_/N_2_ for 8 h, in the dark, for eight hours (from 8 am to 4 pm), while the control plants were kept in the dark under atmospheric conditions (21% ambient O_2_). Each biological replicate consisted on three thalli (*M. polymorpha*), three young plantlets (*P. patens*) or three young shoots (*S. moellendorfii*, *E. hyemale* and *P. vittata*). Hypoxia and submergence treatments for Western blots analysis or qRT-PCR were carried out on *A. thaliana* 7-day-old seedlings, grown *in vitro* conditions, treated with 0.1% v/v O_2_/N_2_ for 2 h, in the dark, or placed in 2 mL Eppendorf tubes filled with deionised water for 1 h, in the dark, respectively. Chemical treatments were performed on *in vitro* Arabidopsis 7-day-old seedlings treated with 50 μM bortezomib (CAS 179324-69-7, Santa Cruz Biotechnology), 100 μM cPTIO (cat. No. C221, Sigma-Aldrich) or DMSO (control), dissolved in deionised water, on an orbital shaker with rotation at 100 rpm, for three and two hours respectively.

### RNA extraction and qRT-PCR analysis

Total RNA was isolated from 80-100 mg of frozen and grinded thirteen-day-old *M. polymorpha* thalli, or *S. moellendorfii*, *E. hyemale* and *P. vittata* young shoots using Spectrum™ Plant Total RNA Kit (Sigma-Aldrich) and GeneJET Plant RNA Purification Kit (Thermo Fisher Scientific), according to the manufacturer’s protocol. For *A. thaliana* 7-day-old seedlings, total RNA was extracted using the RNeasy Plant Mini Kit (Qiagen), following the manufacturer’s instructions. DNase treatments and cDNA synthesis was performed on one microgram of RNA using the Maxima First Strand cDNA Synthesis Kit for RT-qPCR with dsDNase (Thermo Fisher Scientific). Quantitative real-time PCR amplification (qRT-PCR) was carried out for *M. polymorpha*, *S. moellendorfii* and *A. thaliana* RNA samples with the CFX384 Touch Real-Time PCR Detection System (BioRad), using a SYBR® Green PCR Master Mix (Life Technologies). *UBIQUITIN10* (*UBIQ10*, *At4g05320*), *Actin1* (*MpACT1*, *Mapoly0016s0137*) and *6-Phosphogluconate dehydrogenase* (*Sm6PGD*, *98399*) were used as housekeeping genes for *A. thaliana*, *M. polymorpha* and *S. moellendorfii*, respectively. The primer sequences used for the qRT-PCR analysis are listed in **Supplementary Table 36**. Relative quantification of the expression of each gene was performed using the comparative threshold cycle method, as described in (Livak and Schmittgen 2001). Two technical replicates were used for each of the four biological samples and the data are representative of at least two independent experiments giving comparable profiles.

### RNA sequencing analysis

The RNA sequencing reads were first quality trimmed by removal of base calls with a poor Phred score (PHRED Score>30). The reads were then aligned with the Kallisto tool (Bray et al. 2016) to the reference transcriptomes of the corresponding species for *M. polymorpha*, *P. patens*, *S. moellendorffii*, and the *de novo* assembled transcriptomes of *E. hyemale* and *P. vittata*. The *de novo* transcriptome assemblies were performed with the quality trimmed reads and Trinity program using *k*-mer size of 31 and minimal *k*-mer abundance of 2 (Grabherr et al. 2011). Loci/genes where no sample had more than 20 reads were removed from the dataset. Next the libraries were normalized for composition and size with the TMM (trimmed mean of M values) approach (Robinson and Oshlack 2010). To subsequently calculate abundance, fold changes and significance, a negative binomial model was used and tagwise dispersion was estimated with the “edgeR” R package (Robinson and Oshlack 2010). The P-values were adjusted to a false discovery rate of 0.05 with Benjamini-Hochberg correction (Hochberg 1995).

Seven-day-old *A. thaliana* seedlings of Col-0 and *erfVII* mutant were treated with hypoxia (1% v/v O_2_) or control conditions (21% ambient O_2_), in the dark, starting at the end of the day for two and ten hours. Total RNA was extracted using the GeneJET Plant RNA Purification Kit (Thermo Fisher Scientific). Genomic DNA was removed using the AMPure XP system (Beckman Coulter, Beverly, USA). Libraries quality was assed on the Agilent Bioanalyzer 2100 system. The clustering of the index-coded samples was performed on a cBot Cluster Generation System using TruSeq PE Cluster Kit v3-cBot-HS (Illumia) according to the manufacturer’s instructions. The library preparations were sequenced on an Illumina NovaSeq 6000 platform and 150 bp paired-end reads were generated. Paired-end clean reads were aligned to the *A. thaliana* TAIR10 (Ensembl Plants genome ID: 3702) reference genome using Hisat2 v2.0.5. The mapped reads of each sample were assembled by StringTie (v1.3.3b) (Pertea et al. 2015) in a reference-based approach. featureCounts v1.5.0-p3 (Liao, Smyth, and Shi 2014) was used to count the reads numbers mapped to each gene. Fragments Per Kilobase of transcript sequence per Millions base pairs sequenced (FPKM) of each gene was calculated based on the length of the gene and reads count mapped to this gene. Differential expression analysis was performed using the DESeq2 R package (1.20.0). The resulting *p*-values were adjusted using the Benjamini and Hochberg’s approach for controlling the false discovery rate (FDR). Genes were assigned as differentially expressed when the adjusted *p*-value <=0.05 and |log_2_FC|>1 (**Supplementary Table 20**).

### In silico analysis

Gene expression profiles of the differentially expressed genes (DEGs) found in the RNA-seq dataset, selecting log_2_FC>1, log_2_FC<-1 and p-value<0.05, were compared across different experimental conditions by exploring public microarray data based on *A. thaliana* under 1% v/v O_2_ (Licausi et al. 2011) conditions, and on *O. sativa* under anoxic conditions (Narsai et al. 2009) selecting DEGs based on a log_2_FC>1, log_2_FC<-1 and p-value<0.05 (*A. thaliana*) or p-value<0.01 (*O. sativa*).

To identify orthogroups we used 33 species representing key land plant species and the OrthoFinder algorithm (Emms and Kelly 2015). To facilitate the incorporation of *de novo* assembled transcriptomes we favoured a discontiguous Megablast (Altschul et al. 1990) over the default diamond alignment program to make the all-vs-all comparisons. The inflation parameter that controls the granularity of the orthogroup clustering was chosen to maximize the number of conserved orthogroups and set at 1.3.

To identify promoter binding sites we relied on established position-specific scoring matrices (PSSMs) for the HRPE, GCC, MDM, and group I and II R2R3-MYB binding sites (**Supplementary Table 37)**. We used the FIMO tool to scan, genome wide, the 1000 bp upstream and 100 bp downstream of the predicted transcription starts sites. Enrichment of binding motifs was tested using Fisher-exact test.

### SDS-PAGE and immunoblotting

Equal total protein amount (70 μg) were resolved by SDS-PAGE, and were transferred to PVPF membrane using MiniTrans-Blot electrophoretic transfer cell (Bio-Rad). Membranes were probed with anti-HA primary antibody (Sigma-Aldrich; H3663), 1:2000 dilution, followed by HRP-conjugated anti-mouse secondary antibody (Sigma-Aldrich; 12-349), 1:10000 dilution. Immunoblots were developed to films using Pierce™ ECL Western Blotting Substrate (Thermo Fisher Scientific) or with SuperSignal™ West Pico PLUS Chemiluminescent Substrate (Thermo Fisher Scientific) using the iBright CL1500 Imaging System (Thermo Fisher Scientific).

### Confocal imaging

*M. polymorpha* gemmae were used for probe detection after hypoxic treatment (1% v/v O_2_/N_2_) or control conditions (21% ambient O_2_), for six hours, in the dark. For propidium iodide (PI) staining, gemmae were incubated with 10 µg mL-1 of PI solution for five to ten minutes, then rinsed in deionised water and mounted on slide. Imaging was performed using ZEISS LSM 880 microscope (Department of Plant Sciences, University of Oxford), equipped with a 25x objective lens. GFP, PI and chlorophyll were excited at 488nm and emission was collected at 495-560 nm for GFP, 590-635 nm for PI and 650-750 nm.

### Luciferase transactivation assay in protoplasts

*A. thaliana erfVII prt6* or Col-0 (wild-type) mesophyll protoplasts were isolated from leaves of 3-week-old plants and transfected according to (Yoo, Cho, and Sheen 2007). Two micrograms of each plasmid were used to transfect 100 μL of protoplasts suspension. After 16 h of incubation in the dark at 22°C, the firefly (*Photinus pyralis*) luciferase activity was measured and normalized with the sea pansy (*Renilla reniformis*) luciferase signal using the Dual-Luciferase Reporter Assay System (Promega) according to the manufacturer’s instructions. Each sample consisted of five independent replicates.

### Statistical analysis

Statistical analysis were performed using GraphPad Prism version 10. According to the data sets, one-way or two-way ANOVA were conducted, and differences between means were considered significant at *p*-value<0.05. In one-way ANOVA, the multiple comparisons of means were performed with Tukey’s method, whereas in two-way ANOVA, they were performed via the Holm-Sidak method.

## Data availability

All RNA sequencing raw data generated for this study can be accessed at the Sequence Read Archive at the National Centre for Biotechnology Information, under accession numbers PRJNA1128557 and PRJNA1150224.

## Author contributions

M.J.H and F.L conceived the research. L.D.C., HVV, M.J.H, F.L. designed the experiments and interpreted the data; all authors carried out the research; F.L., L.D.C and M.J.H. wrote the manuscript. F.L. agrees to serve as the author responsible for contact and ensures communication. All authors read, contributed to editing and approved the manuscript.

