## Supplementary information for "The role of ERFVIIs as oxygen-sensing transducers in the evolution of land plant response to hypoxia"

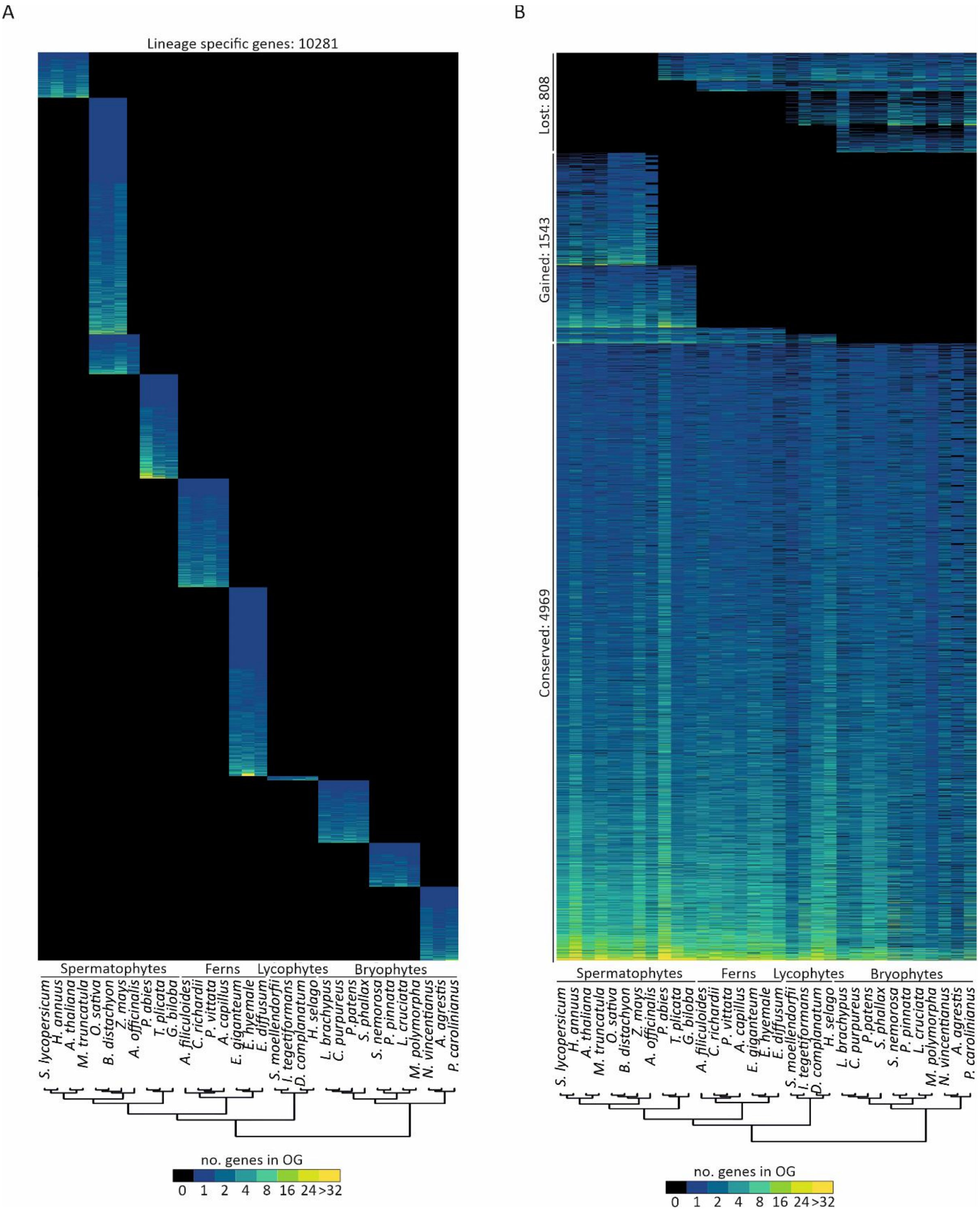


**Supplementary Figure S1**. **Distribution of genes across land plant evolution**. **A**) Hierarchical clustering of lineage specific genes (10281) from 33 land plant species, including those studied in this work: *M. polymorpha*, *P. patens*, *S. moellendorfii*, *E. hyemale*, *P. vittata*, *O. sativa* and *A. thaliana*. **B**) Conserved (4969), gained (1543) and lost (808) orthogroups from 33 land plant species.

A

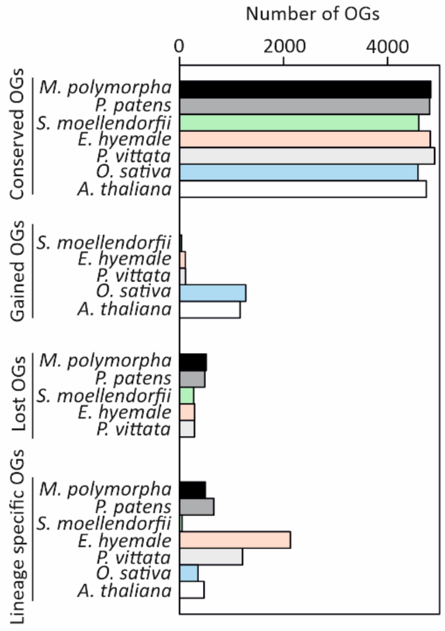


B

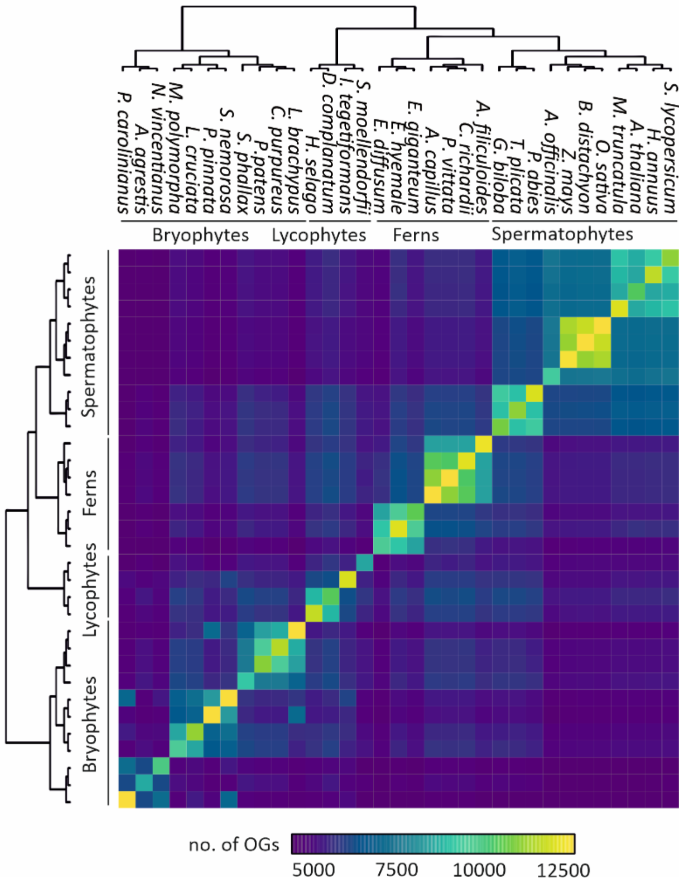


**Supplementary Figure S2**. **Classification of orthogroups across land plant evolution**. **A**) Bar plots depicting the number of conserved, gained, lost and lineage specific orthogroups in *M. polymorpha*, *P. patens*, *S. moellendorfii*, *E. hyemale*, *P. vittata*, *O. sativa* and *A. thaliana*. **B**) The amount of orthogroups shared between pairs of species.

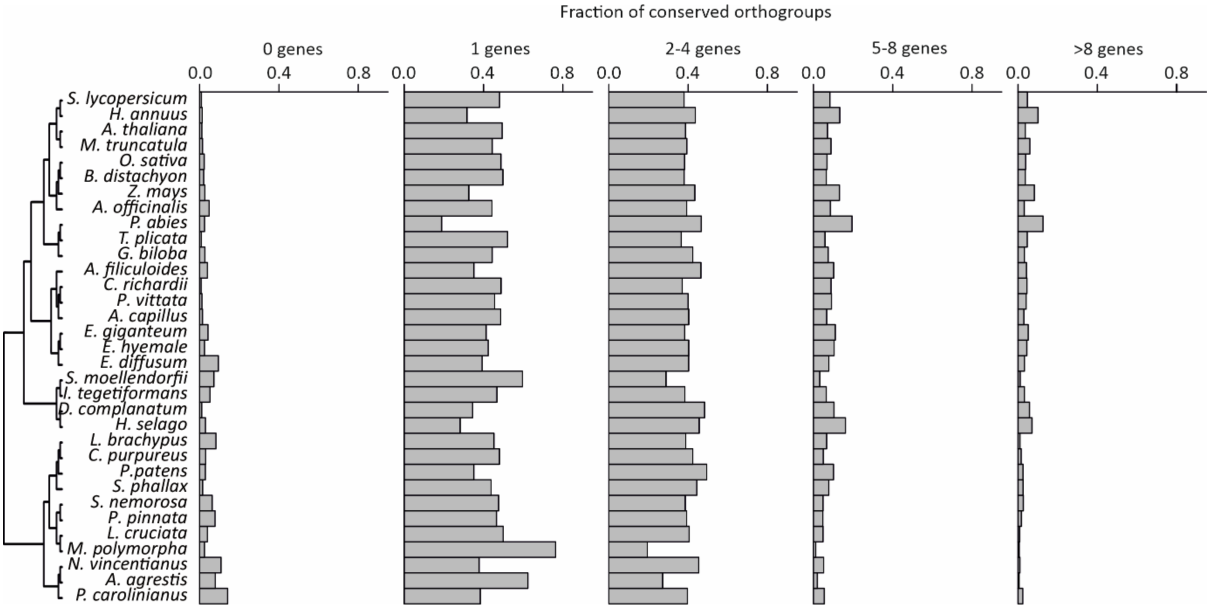


**Supplementary Figure S3. Distribution of number of genes in conserved orthogroups.** Percentage of number of genes, over the total number of genes, included in each of the fraction of conserved orthogroups identified for the 33 land plant species analysed in this study. Fractions of conserved orthogroups are classified according to the number of genes they contain (0, 1, 2-4, 5-8 and more than 8 genes).

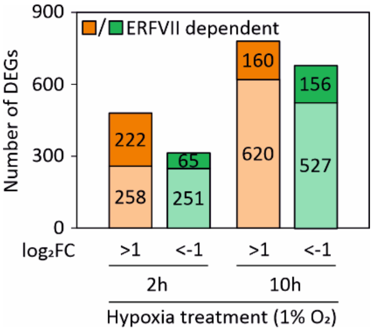


**Supplementary Figure S4**. **Contribution of the ERFVIIs to the hypoxia response in *A. thaliana***. Bar plots representing the number of differentially expressed genes (DEGs) in *A. thaliana* following two or ten hours of dark hypoxia treatment (1% v/v $O_2$) in 7-day-old seedlings of WT (Col-0) plants compared to the *erfVII* mutant. Light orange and light green represent upregulated ($log_2FC>1$, FDR<0.05) or downregulated ($log_2FC<-1$, FDR<0.05) genes, respectively. Dark orange and dark green represent ERFVII-dependent upregulated ($log_2FC>1$, FDR<0.05) or downregulated ($log_2FC<-1$, FDR<0.05) genes, respectively. The exact number of genes for each category is highlighted inside each bar.

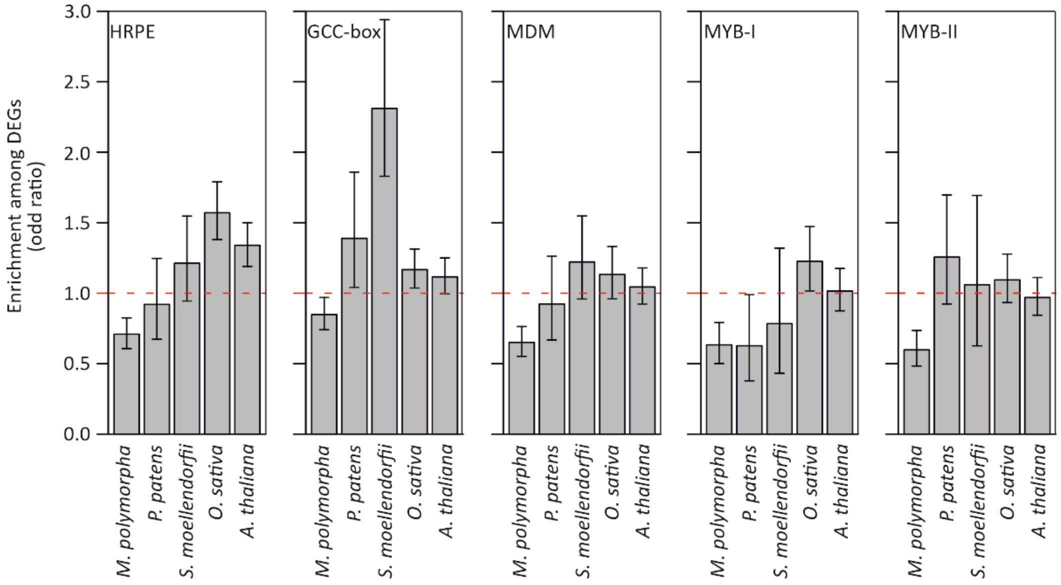


**Supplementary Figure S5**. **Motifs enrichment upon hypoxia in land plants**. Enrichment of *cis*-element motifs (HRPE, GCC-box, MDM, MYB-I and MYB-II) among differentially upregulated genes (P<0.05 and |$log_2$FC|>1). Odds ratio of 1 indicates no enrichment, whereas higher values are enriched. Error bars represent the 95% confidence intervals (Fisher exact test).

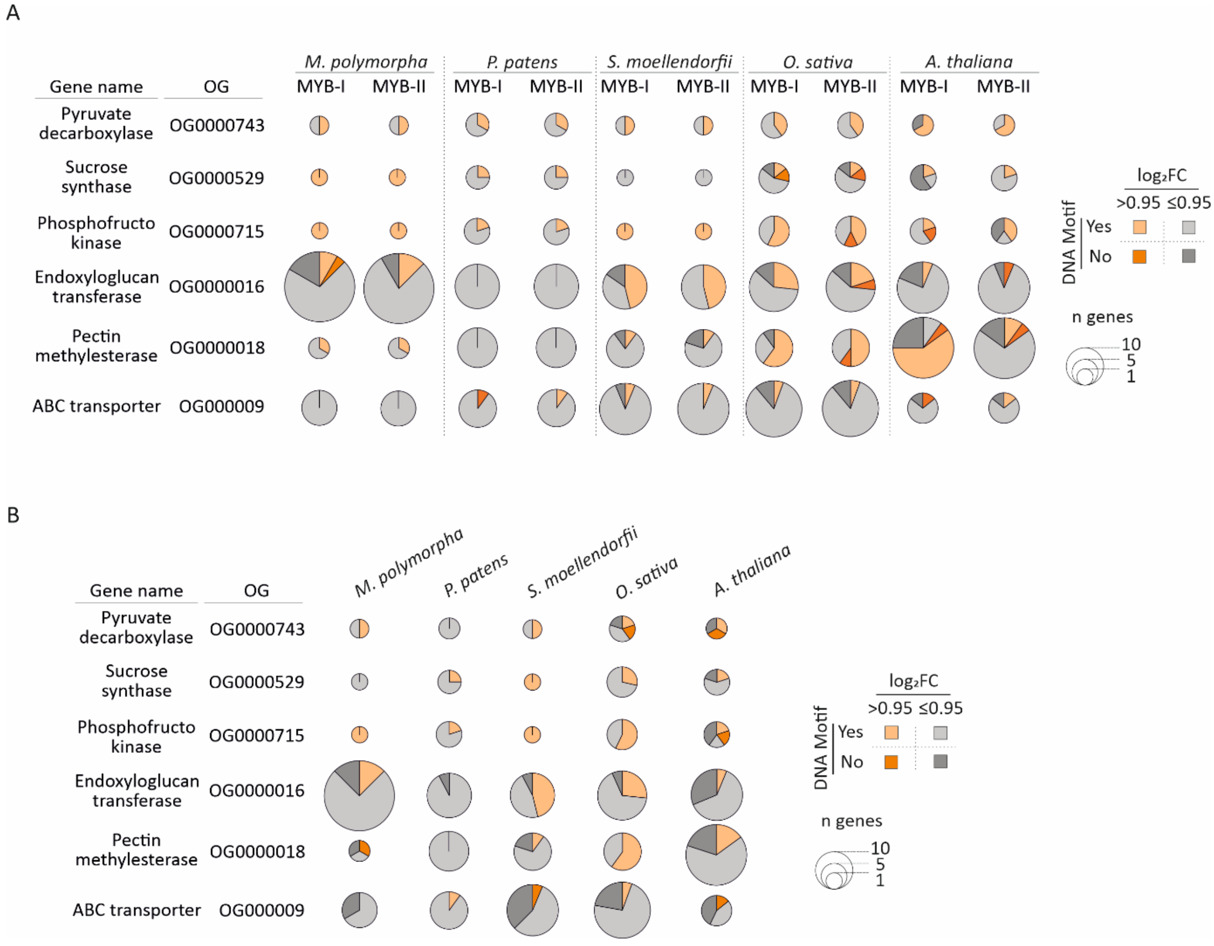

**Supplementary Figure S6**. **Frequency of MYB and NAC *cis*-element motifs in hypoxia-induced promoters across land plants**. **A**) Pie charts representing the occurrence of MYB-binding motifs (MYB-I and MYB-II) or **B**) NAC-binding motif (MDM) in promoters of upregulated (dark orange, $log_2FC>0.95$) and downregulated (dark grey, $log_2FC \leq 0.95$) genes belonging to the seven hypoxia-induced and conserved orthogroups in *M. polymorpha*, *P. patens*, *S. moellendorfii*, *E. hyemale*, *P. vittata*, *O. sativa* and *A. thaliana*. The size of each pie chart is proportional to the number of genes belonging to the orthogroups for each individual species, with total upregulated genes in light orange ($log_2FC>0.95$) and total downregulated genes in light grey ($log_2FC \leq 0.95$).

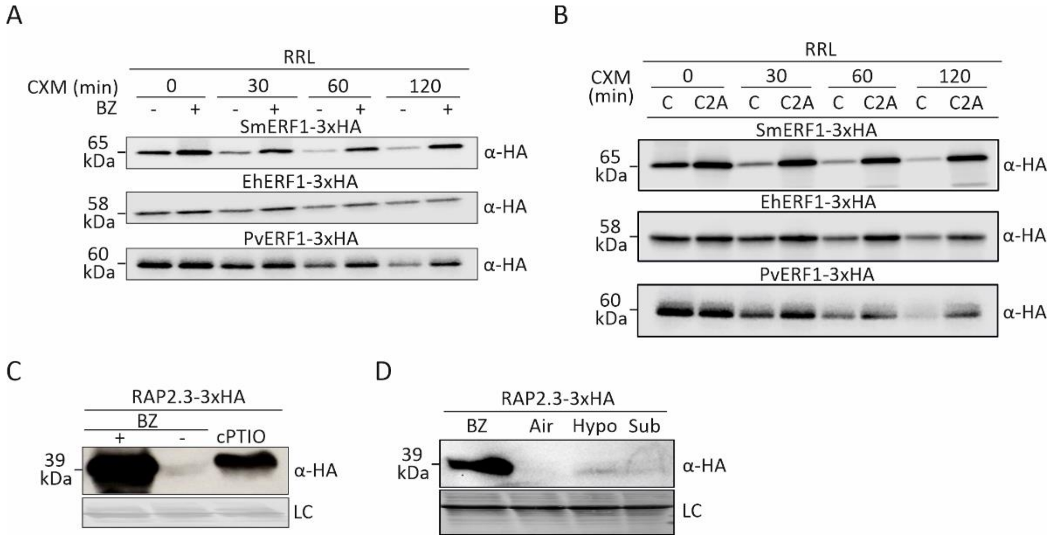


**Supplementary Figure S7**. **Tracheophyte ERFVIIs regulation through the PCO branch of the PRT6 N-degron pathway**. **A**) Western blot analysis of SmERF1$^{3xHA}$, EhERF1$^{3xHA}$ and PvERF1$^{3xHA}$ abundance in rabbit reticulocyte lysate (RRL) in control (DMSO) or bortezomib (BZ) conditions during a time course (0, 30, 60 and 120 min) after cycloheximide (CXM) addition. **B**) Western blot analysis of Cys2 compared to C2A SmERF1$^{3xHA}$, EhERF1$^{3xHA}$ and PvERF1$^{3xHA}$ abundance in RRL during a time course (0, 30, 60 and 120 min) after cycloheximide (CXM) treatment. **C**) Western blot analysis of RAP2.3$^{3xHA}$ abundance in control (DMSO), bortezomib (BZ) or NO-scavenger (cPTIO) treated *erfVII* mutant seedlings. **D**) Western blot analysis of RAP2.3$^{3xHA}$ abundance in bortezomib (BZ), air (21% v/v $O_2$), hypoxia (2h, 0.1% v/v $O_2$) or submergence (1h) treated *erfVII* mutant seedlings. Loading control (LC) shown corresponds to a Ponceau staining of total loaded proteins on Western blots.

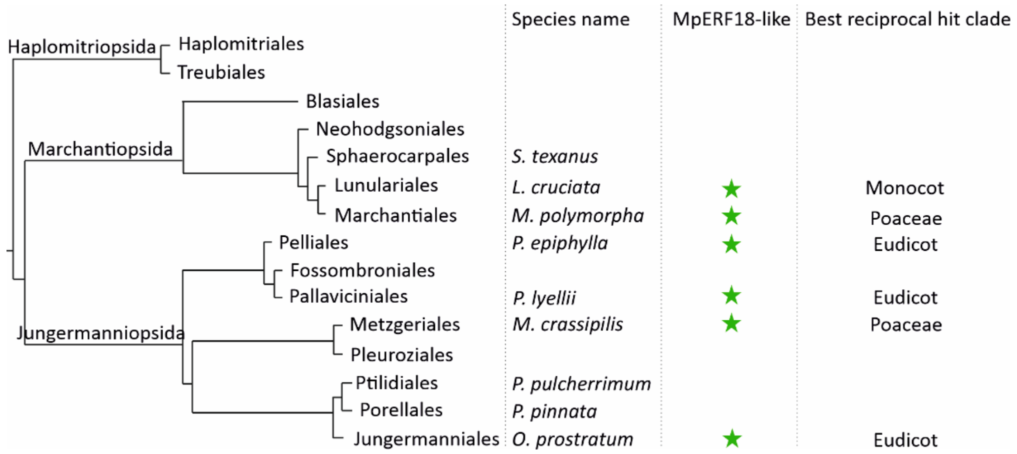

**Supplementary Figure S8**. **Occurrence of *M. polymorpha* MpERF18-like proteins in liverworts**. tblastn algorithm search of the 1KP database using *M. polymorpha* MpERF18 (Mapoly0092s0058) protein sequence as query. The presence of orthologues of MpERF18 are marked with a green star, together with the species name where they have been identified. For each MpERF18 orthologs found, their best reciprocal hit clade is as well reported.

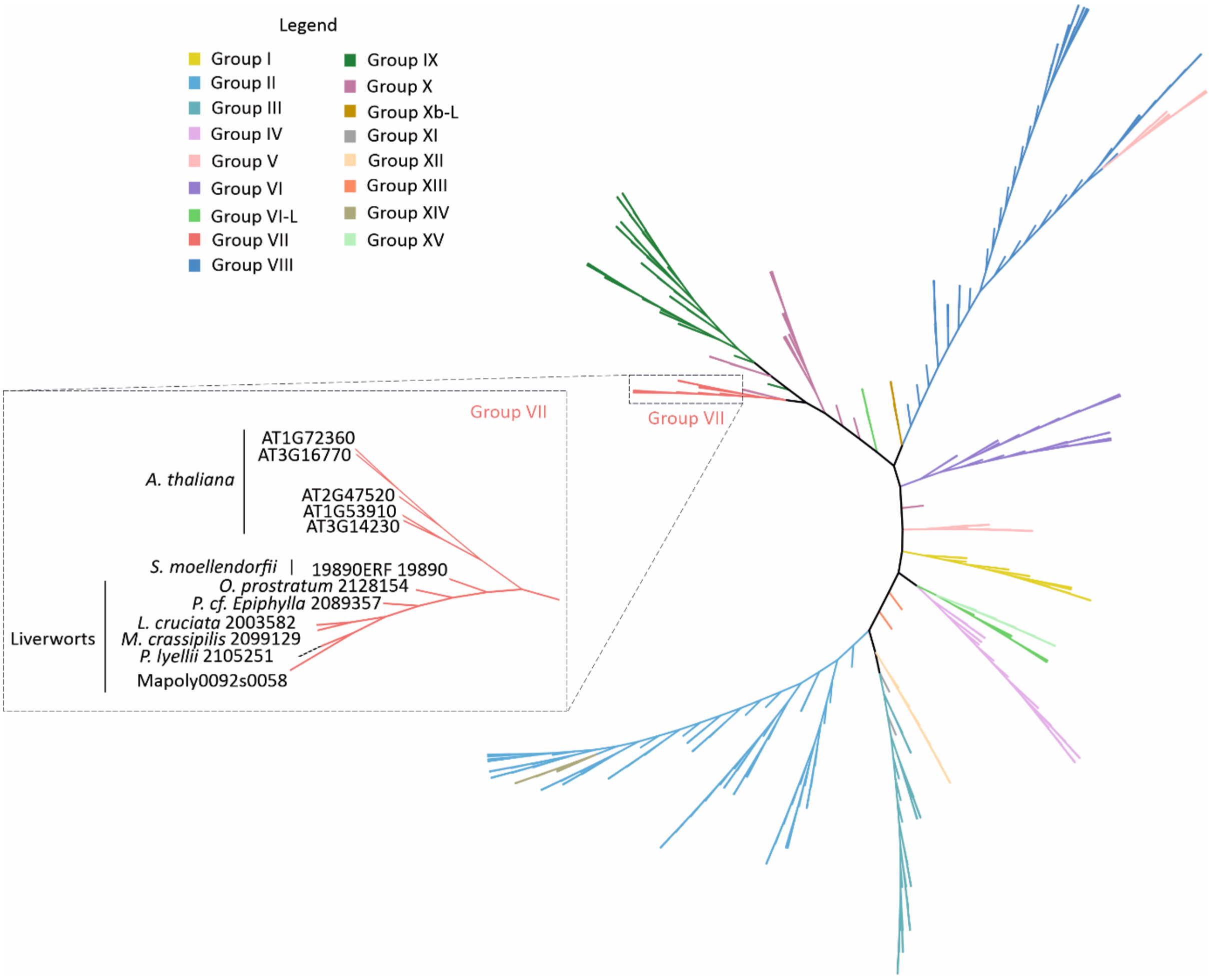


**Supplementary Figure S9**. **Phylogenetic relationship between ERF proteins in bryophytes, *S. moellendorfii* and *A. thaliana*.** Unrooted phylogenetic tree of ERF proteins from bryophytes (*A. punctatus*, *M. polymorpha* and *P. patens*), lycophyte (*S. moellendorfii*) and angiosperm (*A. thaliana*), including orthologous ERFVII-like proteins of MpERF18 retrieved from other groups of liverworts, which are shown in **Supplementary Fig. S8** and listed in **Supplementary Table 34**. ERFs classification is based on (Nakano et al. 2006). In the dashed box is a magnification of the Group VII branch, where the names of the sequences at each node and the species or clade to which they belong are highlighted.

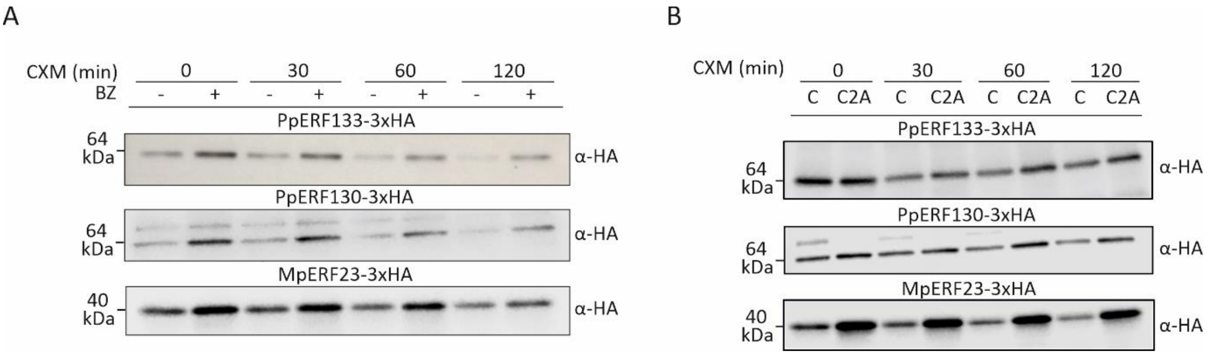


**Supplementary Figure S10**. **Bryophyte ERF regulation through the N-degron pathway. A**) Western blot analysis of ERF Cys2 proteins PpERF133$^{3xHA}$, PpERF130$^{3xHA}$ and MpERF23$^{3xHA}$ abundance in rabbit reticulocyte lysate (RRL) in control (DMSO) or bortezomib (BZ) conditions during a time course (0, 30, 60 and 120 min) after cycloheximide (CXM) treatment. **B**) Western blot analysis of Cys2 compared to C2A PpERF133$^{3xHA}$, PpERF130$^{3xHA}$ and MpERF23$^{3xHA}$ abundance in RRL during a time course (0, 30, 60 and 120 min) after cycloheximide (CXM) treatment.

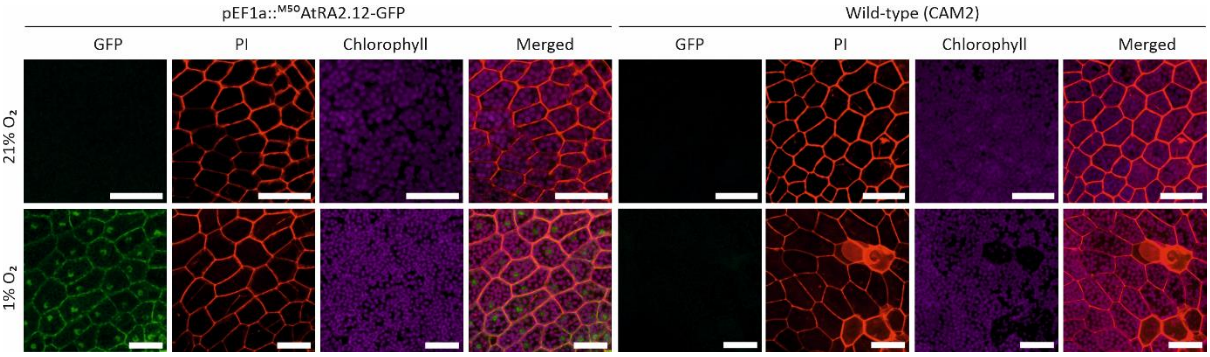


**Supplementary Figure S11**. ***A. thaliana*** **RAP2.12 stability depends on hypoxia in *M. polymorpha***. Confocal images of the subcellular localization of a reporter consisting of the first 50 aa of AtRAP2.12 fused to Citrine, overexpressed in *M. polymorpha* 13-day-old thalli in air (21% ambient $O_2$) and hypoxia (1% v/v $O_2$), compared to wild-type (CAM2) plants.

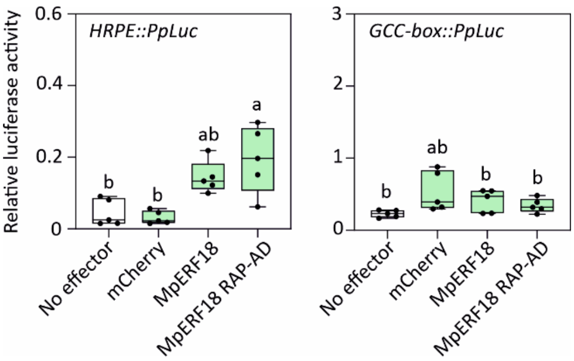


**Supplementary Figure S12**. **MpERF18 transactivation efficiency on hypoxia-related promoters**. Effect of *M. polymorpha* ERFVII-like (MpERF18) wild type or fused at the C-terminal to RAP2.12 activation domain (MpERF18 RAP-AD) on the synthetic promoters HRPE (left) and GCC-box (right), measured as relative firefly luciferase activity in *A. thaliana* wild-type mesophyll protoplasts. Data are presented as means ± SD (n=5). Different letters indicate statistically different averages as assessed by one-way ANOVA followed by the Tukey's posthoc test ($P < 0.05$).

**Supplementary Table 1.** RNAseq analysis of *Marchantia polymorpha* in control (21% v/v $O_2$) and hypoxia (1% v/v $O_2$), for eight hours in the dark.

**Supplementary Table 2**. RNAseq analysis of *Physcomitrium patens* in control (21% v/v $O_2$) and hypoxia (1% v/v $O_2$), for eight hours in the dark.

**Supplementary Table 3**. RNAseq analysis of *Selaginella moellendorfii* in control (21% v/v $O_2$) and hypoxia (1% v/v $O_2$), for eight hours in the dark.

**Supplementary Table 4**. RNAseq analysis of *Equisetum hyemale* in control (21% v/v $O_2$) and hypoxia (1% v/v $O_2$), for eight hours in the dark.

**Supplementary Table 5**. RNAseq analysis of *Pteris vittata* in control (21% v/v $O_2$) and hypoxia (1% v/v $O_2$), for eight hours in the dark.

**Supplementary Table 6.** List of *M. polymorpha* genes associated to each orthogroup defined according the OrthoFinder method (Emms and Kelly 2015), alongside with the corresponding $log_2FC$ and fold discovery rate (FDR), in hypoxia.

**Supplementary Table 7**. List of *P. patens* genes associated to each orthogroup defined according the OrthoFinder method (Emms and Kelly 2015), alongside with the corresponding $log_2FC$ and fold discovery rate (FDR), in hypoxia.

**Supplementary Table 8**. List of *S. moellendorfii* genes associated to each orthogroup defined according the OrthoFinder method (Emms and Kelly 2015), alongside with the corresponding $log_2FC$ and fold discovery rate (FDR), in hypoxia.

**Supplementary Table 9**. List of *E. hyemale* genes associated to each orthogroup defined according the OrthoFinder method (Emms and Kelly 2015), alongside with the corresponding $log_2FC$ and fold discovery rate (FDR), in hypoxia.

**Supplementary Table 10**. List of *P. vittata* genes associated to each orthogroup defined according the OrthoFinder method (Emms and Kelly 2015), alongside with the corresponding $log_2FC$ and fold discovery rate (FDR), in hypoxia.

**Supplementary Table 11**. List of *O. sativa* genes associated to each orthogroup defined according the OrthoFinder method (Emms and Kelly 2015), alongside with the corresponding $log_2FC$ and fold discovery rate (FDR), in hypoxia.

**Supplementary Table 12**. List of *A. thaliana* genes associated to each orthogroup defined according the OrthoFinder method (Emms and Kelly 2015), and the corresponding $log_2FC$, along with the fold discovery rate (FDR), in hypoxia.

**Supplementary Table 13**. Number of orthogroups identified for each land plant group analysed in this study. Orthogroups are classified in 'conserved', 'gained', 'lost' and 'lineage specific'.

**Supplementary Table 14**. Number of genes that constitute the hierarchical clustering of conserved orthogroups from 33 species of land plants, including those studied in this work.

**Supplementary Table 15**. Percentage of number of genes, over the total number of genes, included in each of the fraction of conserved orthogroups identified for the 33 land plant species analysed in this study.

**Supplementary Table 16**. Number of orthogroups containing exclusively upregulated (log2FC>1) or downregulated (log2FC<-1) genes or simultaneously up- and down- regulated (-1<log2FC>1) genes under hypoxia. Gene cutoff 0 defines the proportion of genes in an orthogroup that need to be differentially expressed for the OG to be considered regulated.

| Species name | $Log_2FC>1$ | -1<Log2FC>1 | $Log_2FC<-1$ |
|---|---|---|---|
| *M. polymorpha* | 263 | 7 | 283 |
| *P. patens* | 28 | 1 | 65 |
| *S. moellendorfii* | 11 | 0 | 148 |
| *E. hyemale* | 92 | 25 | 375 |
| *P. vittata* | 86 | 7 | 639 |
| *O. sativa* | 908 | 56 | 464 |
| *A. thaliana* | 541 | 43 | 492 |

**Supplementary Table 17**. $log_2FC<-1$ (Padj. <0.05) of conserved hypoxia-regulated orthogroups in the land plant species considered in this study. Orthogroups are classified according to the molecular function of the orthologous *A. thaliana* gene.

**Supplementary Table 18**. $log_2FC>1$ (Padj. <0.05) of conserved hypoxia-regulated orthogroups in the land plant species considered in this study. Orthogroups are classified according to the molecular function of the orthologous *A. thaliana* gene.

**Supplementary Table 19**. List of hypoxia-induced and repressed conserved orthogroups in which the corresponding A. thaliana protein belongs to the 'enzyme' class.

**Supplementary Table 20**. RNAseq analysis of *Arabidopsis thaliana* wild-tpe (Col-0) and *erfVII* pentuple mutant after hypoxia treatment time course (2 h and 10 h of 1% v/v O2 in the dark).

**Supplementary Table 21**. HRPE motif enrichment analysis in promoters of orthogroups, divided by species.

**Supplementary Table 22**. GCC-box motif enrichment analysis in promoters of orthogroups, divided by species.

**Supplementary Table 23**. MDM motif enrichment analysis in promoters of orthogroups, divided by species.

**Supplementary Table 24**. MYB-I motif enrichment analysis in promoters of orthogroups, divided by species.

**Supplementary Table 25**. MYB-II motif enrichment analysis in promoters of orthogroups, divided by species.

**Supplementary Table 26**. HRPE motif enrichment on promoter regions of conserved hypoxia-induced orthogroups.

**Supplementary Table 27**. GCC-box motif enrichment on promoter regions of conserved hypoxia-induced orthogroups.

**Supplementary Table 28**. MDM motif enrichment on promoter regions of conserved hypoxia-induced orthogroups.

**Supplementary Table 29**. MYB-I motif enrichment on promoter regions of conserved hypoxia-induced orthogroups.

**Supplementary Table 30**. MYB-II motif enrichment on promoter regions of conserved hypoxia-induced orthogroups.

**Supplementary Table 31**. List or protein sequences from different species of lycophytes, eusporangiate ferns, leptosporangiate ferns, gymnosperms and angiosperms, orthologous of *Arabidopsis thaliana* RAP2.12, used to build the phylogenetic tree of ERFVII transcription factors family in tracheophytes.

**Supplementary Table 32.** List of ERF transcription factors from *Marchantia polymorpha*, *Physcomitrium patens*, *Anthoceros agrestis*, *Selaginella moellendorfii* and *A. thaliana* used to build the phylogenetic tree shown in **Fig. 4**.

**Supplementary Table 33**. List of ERF transcription factors from *Chlamydomonas reinhardtii* and *Klebsormidium flaccidum*.

**Supplementary Table 34.** List of the best *Marchantia polymorpha* ERFVII (Mapoly0092s0058, MpERF18) orthologs retrieved in specific classes and orders of liverworts.

**Supplementary Table 35**. List of ERF transcription factor proteins starting with a cysteine in second position in different species of bryophytes.

**Supplementary Table 36**. List of forward and reverse primer sequences (5'-3') used for various cloning and RT-qPCR analysis in this study.

**Supplementary Table 37**. Overview over Position-specific scoring matrices (PSSMs) for MYB-I and MYB-II *cis*-elements identified in this study. Matrices have been created using MEME version 5.5.7.

| MYB I | | | | MYB II | | | |
|---|---|---|---|---|---|---|---|
| a | c | g | t | a | c | g | t |
| 0.5 | 0 | 0.17 | 0.33 | 0.25 | 0 | 0.75 | 0 |
| 0.33 | 0.17 | 0.17 | 0.33 | 0.125 | 0.125 | 0.5 | 0.25 |
| 0.33 | 0.5 | 0 | 0.17 | 0.1 | 0.1 | 0 | 0.8 |
| 0.67 | 0 | 0.17 | 0.17 | 0.6 | 0 | 0.2 | 0.2 |
| 0 | 0 | 1 | 0 | 0 | 0 | 1 | 0 |
| 0 | 0 | 0 | 1 | 0 | 0 | 0.2 | 0.8 |
| 0 | 0 | 0 | 1 | 0 | 0 | 0 | 1 |
| 0.5 | 0 | 0.5 | 0 | 0.5 | 0 | 0.5 | 0 |

**Supplementary File S1**. Multi alignment between protein sequences identified in *M. polymorpha*, *S. moellendorfii*, *E. hyemale*, *P. vittata*, *O. sativa* and *A. thaliana* belonging to the OG0000005 orthogroup.

**Supplementary File S2**. Multi alignment between protein sequences identified in *M. polymorpha*, *S. moellendorfii*, *E. hyemale*, *O. sativa* and *A. thaliana* belonging to the OG0000004 orthogroup.

**Supplementary File S3**. Multi alignment between protein sequences identified in *M. polymorpha*, *S. moellendorfii*, *E. hyemale*, *O. sativa* and *A. thaliana* belonging to the OG0007434 orthogroup.

**Supplementary File S4**. Multi alignment between protein sequences identified in *M. polymorpha*, *P. patens*, *S. moellendorfii*, *E. hyemale*, *O. sativa* and *A. thaliana* belonging to the OG0000715 orthogroup.

**Supplementary File S5**. Multi alignment between protein sequences identified in *M. polymorpha*, *P. patens*, *S. moellendorfii*, *E. hyemale*, *P. vittata*, *O. sativa* and *A. thaliana* belonging to the OG0000529 orthogroup.

**Supplementary File S6**. Multi alignment between protein sequences identified in *M. polymorpha*, *S. moellendorfii*, *E. hyemale*, *P. vittata*, *O. sativa* and *A. thaliana* belonging to the OG0000018 orthogroup.

**Supplementary File S7**. Multi alignment between protein sequences identified in *M. polymorpha*, *S. moellendorfii*, *E. hyemale*, *P. vittata* and *O. sativa* belonging to the OG0000016 orthogroup.

**Supplementary File S8**. Multi alignment between protein sequences identified in *P. patens*, *S. moellendorfii*, *E. hyemale*, *P. vittata*, *O. sativa* and *A. thaliana* belonging to the OG0000009 orthogroup.

**Supplementary File S9**. Multi alignment between *A. thaliana* five ERFVII protein sequences and the best ERF blast hits retrieved in bryophytes using AtRAP2.12 as query.

**Supplementary File S10**. Multi alignment between *A. thaliana* five ERFVII protein sequences and the best ERFVII orthologs identified in *Equisetum hyemale* and *Equisetum diffusum*, using AtRAP2.12 as query.
